# Re-programming of GM-CSF-dependent alveolar macrophages through GSK3 activity modulation

**DOI:** 10.1101/2024.08.20.608749

**Authors:** Israel Ríos, Cristina Herrero, Mónica Torres-Torresano, Baltasar López-Navarro, María Teresa Schiaffino, Francisco Díaz-Crespo, Alicia Nieto-Valle, Rafael Samaniego, Yolanda Sierra-Palomares, Eduardo Oliver, Fernando Revuelta-Salgado, Ricardo García-Luján, Paloma Sánchez-Mateos, Rafael Delgado, Amaya Puig-Kröger, Ángel L. Corbí

**Author notes:** Corresponding authors: Dr. Ángel L. Corbí. Centro de Investigaciones Biológicas “Margarita Salas”, CSIC. Ramiro de Maeztu, 9. Madrid 28040; Dr. Amaya Puig-Kröger. Unidad de Inmuno-Metabolismo e Inflamación, Instituto de Investigación Sanitaria Gregorio Marañón (IiSGM), Madrid, Spain. APK and ALC contributed equally to this work.

## Abstract

Monocyte-derived macrophages recruited into inflamed tissues can acquire an array of functional states depending on the extracellular environment. Since the anti-inflammatory/pro-fibrotic macrophage profile is determined by MAFB, whose activity/protein levels are regulated by GSK3, we addressed the macrophage re-programming potential of GSK3 modulation. GM-CSF-dependent (GM-MØ) and M-CSF-dependent monocyte-derived macrophages (M-MØ) exhibited distinct levels of inactive GSK3, and inhibiting GSK3 in GM-MØ led to acquisition of transcriptional, phenotypic and functional properties characteristic of M-MØ (enhanced expression of IL-10 and monocyte-recruiting factors, and higher efferocytosis). These re-programming effects were also observed upon GSK3α/β knockdown, and through GSK3 inhibition in *ex vivo* isolated human alveolar macrophages (AMØ). Notably, GSK3 downmodulation potentiated the transcriptional signature of Interstitial Macrophages (IMØ) while suppressed the AMØ-specific gene profile. Indeed, heightened levels of inactive GSK3 and MAFB-dependent proteins were observed in severe COVID-19 patients lung macrophages, highlighting the GSK3-MAFB axis as a therapeutic target for macrophage re-programming.

## INTRODUCTION

Macrophages exhibit considerable functional plasticity during inflammatory responses, transitioning from pro-inflammatory activities to tissue repair and inflammation resolution ^1,2^. This functional versatility of macrophages is intrincately tied to factors such as their ontogeny (fetal origin vs. monocyte-derived) and the specific tissue and extracellular environment ^3–5^. Notably, monocyte-derived macrophages are oppositely instructed by M-CSF or GM-CSF ^6–12^. GM-CSF directs monocyte-derived macrophages (GM-MØ) towards heightened pro-inflammatory activity ^10,13–15^ and the acquisition of the lung alveolar macrophage phenotype and gene profile ^16^. In fact, GM-CSF is essential for generating lung alveolar macrophages ^17^, crucial for surfactant homeostasis, whereas deficient GM-CSF signaling leads to pulmonary alveolar proteinosis ^18^, a condition marked by defective surfactant clearance and disruption of pulmonary homeostasis due to alveolar macrophage dysfunction ^19^. Conversely, M-CSF gives rise to anti-inflammatory, pro-resolving, and immunosuppressive monocyte-derived macrophages (M-MØ) ^1,2,20^ but promotes the development of pro-fibrotic pathogenic monocyte-derived macrophages in lung pathologies like idiopathic pulmonary fibrosis (IPF) ^21^ and severe COVID-19 ^22^. The pathological relevance of GM-MØ and M-MØ subsets is evident in severe COVID-19, with a huge increase in monocyte-derived M-MØ and a reduction of tissue-resident GM-MØ-like lung macrophages ^23–25^.

Despite their crucial role in maintaining tissue homeostasis, deregulated macrophage functional specialization contributes to various human diseases, including tumors and chronic inflammatory pathologies ^26^. Consequently, macrophage re-programming has emerged as a promising therapeutic approach ^26^. However, achieving this necessitates a profound understanding of the transcriptional and signaling mechanisms governing the pro- and anti-inflammatory nature of macrophages. In this regard, recent findings indicate that the homeostatic and reparative transcriptional profile of human M-MØ is orchestrated by MAF and MAFB ^27–29^, closely related transcription factors that regulate stemness and self-renewal in the mouse haematopoietic lineage ^30–35^. In fact, the distinctive expression of MAFB in M-MØ, influencing IL-10 production ^27,36^, also mediates macrophage re-programming induced by methotrexate ^37^ and LXR ligands ^38^.

Given the pivotal role of MAFB/MAF in human macrophage functional specification, and considering the regulation of large MAF factors through GSK3-mediated phosphorylation-induced proteasomal degradation ^30^, we hypothesized that modulating GSK3 activity could offer a viable avenue for re-programming monocyte-derived macrophages. Our findings reveal significant differences in inhibitory phosphorylation of GSK3β (Ser^9^-GSK3β) and GSK3α (Ser^21^-GSK3α) between M-MØ and GM-MØ, and that GSK3 inhibition (using CHIR-99021) or knock-down (using siRNA) in GM-MØ prompts the adoption of transcriptional, phenotypic, and functional characteristics resembling those of M-MØ. This re-programming effect was manifested in elevated expression of IL-10, monocyte-recruiting chemokines, pro-fibrotic factors, as well as in enhanced phagocytic capability, all correlating with increased MAFB expression. The re-programming action of GSK3 inhibition was also observed in control monocytes and in monocytes exposed to a pathological pro-inflammatory environment. Importantly, GSK3 inhibition in *ex vivo* isolated human Alveolar Macrophages (AMØ), led to the loss of the AMØ gene signature and the acquisition of the gene profile that characterizes Interstitial Macrophages (IMØ). Since pathogenic lung macrophages in severe COVID-19 exhibit high levels of inactive GSK3 and heightened levels of MAFB and MAFB-dependent proteins, our results underscore the potent macrophage re-programming impact of GSK3 inhibition, and highlight the GSK3-MAFB axis as a promising therapeutic target for macrophage re-programming.

## EXPERIMENTAL PROCEDURES

### Generation of human monocyte-derived macrophages *in vitro* and treatments

Human Peripheral Blood Mononuclear Cells (PBMCs) were isolated from buffy coats from anonymous healthy donors over a Lymphoprep (Nycomed Pharma) gradient according to standard procedures. Monocytes were purified from PBMC by magnetic cell sorting using anti-CD14 microbeads (Miltenyi Biotec). Monocytes (>95% CD14^+^ cells) were cultured at 0.5 × 10^6^ cells/ml in Roswell Park Memorial Institute (RPMI 1640, Gibco) medium supplemented with 10% fetal bovine serum (FBS, Biowest) (complete medium) for 7 days in the presence of 1000 U/ml GM-CSF or 10 ng/ml M-CSF (ImmunoTools) to generate GM-CSF-polarized macrophages (GM-MØ) or M-CSF-polarized macrophages (M-MØ), respectively ^27^. Cytokines were added every two days and cells were maintained at 37°C in a humidified atmosphere with 5% CO_2_ and 21% O_2_. Alveolar macrophages (AMØ) were obtained from remains of bronchoalveolar lavage (BAL) of patients undergoing bronchoscopy for diagnostic purposes under a protocol approved by the Internal Review Board of Instituto de Investigación Hospital 12 de Octubre (Reference TP24/0183). BAL procedure was performed with a flexible bronchoscope and a total volume of 150 ml of sterile isotonic saline solution at 37°C. BAL fluid fractions were maintained at 4°C and cellular debris removed using a 40 μm cell strainer. BAL cells were washed with PBS, centrifuged and resuspended in complete medium containing 100 U/mL penicillin and 100 μg/mL streptomycin (#15140-122,Gibco), 50 μg/ml gentamicin (#G1397, Sigma-Aldrich), and 2.5 μg/ml amphotericin B (#A2942, Sigma-Aldrich). The cells were seeded at 6–8 × 10^5^ cells per well in 12-well plates for 1 h and washed extensively to remove non-adherent cells. Finally, 2 ml of complete medium with antibiotics was added to each well and the adherent cells incubated for 16–18 h before treatments. More than 95% of adherent BAL cells were identified as AMØ according to morphology and phenotypic analysis. When indicated, monocytes, GM-MØ or AMØ were exposed to the GSK3 inhibitors CHIR99021 (10 μM), SB-216763 (10 μM) or LiCl (10 mM), using DMSO as control. For macrophage activation, cells were treated with either 100 ng/ml CL264 (InvivoGen), 10 ng/ml *E. coli* 0111:B4 lipopolysaccharide (LPS-EB Ultrapure, Invivogen), TNF (20 ng/ml, ImmunoTools) or IFNγ (5 ng/ml, ImmunoTools). Human cytokine production was measured in M-MØ culture supernatants using commercial ELISA [CCL2 (BD Biosciences), IL-10, CCL18, LGMN, SPP1 (R&D Systems) and activin A (BD Biosciences)] and following the procedures supplied by the manufacturers.

### siRNA transfection

GM-MØ (1 × 10^6^ cells) were transfected with a human *GSK3A*-specific and/or *GSK3B*-specific siRNA (50 nM) (Dharmacon) using HiPerFect (Qiagen). Silencer Select Negative Control No. 2 siRNA (siCtrl, 50 nM) (Dharmacon) was used as negative control siRNA. Six hours after transfection, cells were either allowed to recover from transfection in complete medium (66 hours), and lysed. Knockdown of GSK3α/β was confirmed by Western blot.

### Quantitative real-time RT-PCR (qRT-PCR)

Total RNA was extracted using the total RNA and protein isolation kit (Macherey-Nagel). RNA samples were reverse-transcribed with High-Capacity cDNA Reverse Transcription reagents kit (Applied Biosystems) according to the manufacturer’s protocol. Real-time quantitative PCR was performed with LightCycler® 480 Probes Master (Roche Life Sciences) and Taqman probes on a standard plate in a Light Cycler® 480 instrument (Roche Diagnostics). Gene-specific oligonucleotides (*IL10*: Forward, 5’-TCACTCATGGCTTTGTAGATGC-3’, and Reverse, 5’-GTGGAGCAGGTGAAGAATGC-3’; *LGMN*: Forward, 5’-GAACACCAATGATCTGGAGGA-3’, and Reverse, 5’-GGAGACGATCTTACGCACTGA-3’) were designed using the Universal ProbeLibrary software (Roche Life Sciences). Results were normalized to the expression level of the endogenous references genes (*TBP*, *HPRT1*) and quantified using the ΔΔCT (cycle threshold) method.

### Western blot

Cell lysates were subjected to SDS-PAGE (50 μg unless indicated otherwise) and transferred onto an Immobilon-P polyvinylidene difluoride membrane (PVDF; Millipore). After blocking the unoccupied sites with 5% non-fat milk diluted in Tris-Buffered Saline plus Tween 20 (TBS-T), protein detection was carried out with antibodies against MAFB (HPA005653, Sigma Aldrich), total GSK3α (#4337, Cell Signaling Technology), total GSK3β (#27C10, Cell Signaling Technology), total GSK3 (#5676, Cell Signaling Technology), p-Ser^9^-GSK3β (#5558, Cell Signaling Technology), p-Ser^21^-GSK3α (#9316, Cell Signaling Technology), p-Tyr^279^/P-Tyr^216^– GSK3α/β (#05-413, Sigma Aldrich), CD163 (#MCA1853, Bio-Rad), MAF (sc-518062, Santa Cruz Biotechnology), IL7R (sc-514445, Santa Cruz Biotechnology) and FOLR2 (clone 94b, kindly provided by Dr. Takami Matsuyama) ^39^, and vinculin (#V9131, Sigma-Aldrich) or GAPDH (sc-32233, Santa Cruz Biotechnology) as protein loading controls. Quimioluminiscence was detected in a Chemidoc Imaging system (BioRad) using SuperSignal™ West Femto (ThermoFisher Scientific).

### RNA-sequencing and data analysis

RNA isolated from GM-MØ exposed to DMSO or the GSK3β inhibitor CHIR-99021 (10 μM) for 48h, monocytes exposed to CHIR99021 or DMSO, *ex vivo* isolated AMØ exposed to CHIR99021 or DMSO, or GM-MØ transfected with siRNA specific for *GSK3A* and/or *GSK3B*, or a control siRNA, were subjected to sequencing on a BGISEQ-500 platform (https://www.bgitechsolutions.com). RNAseq data were deposited in the Gene Expression Omnibus (http://www.ncbi.nlm.nih.gov/geo/) under accession GSE256538 (monocytes exposed to CHIR99021 or DMSO), GSE256208 (GM-MØ exposed to CHIR99021 or DMSO), GSE262463 (AMØ exposed to CHIR99021 or DMSO) or GSE266236 (GM-MØ transfected with siRNA specific for *GSK3A*, *GSK3B* or a control siRNA). Low quality reads and reads with adaptors or unknown bases were filtered to get the clean reads. Sequences were mapped to GRCh38 genome using HISAT2 ^40^ or Bowtie2 ^41^, and clean reads for each gene were calculated using htseq-count ^42^ and the RSEM software package ^43^. Differential gene expression was assessed by using the R-package DESeq2. Differentially expressed genes were analysed for annotated gene sets enrichment using ENRICHR (http://amp.pharm.mssm.edu/Enrichr/) ^44,45^, and enrichment terms considered significant with a Benjamini-Hochberg-adjusted p value <0.05. For gene set enrichment analysis (GSEA) (http://software.broadinstitute.org/gsea/index.jsp) ^46^, gene sets available at the website, as well as gene sets generated from publicly available transcriptional studies (https://www.ncbi.nlm.nih.gov/gds), were used. The datasets used throughout the manuscript (either reported here for the first time or previously published) are listed and described in Table S1.

### Re-analysis of single cell RNA-seq data

Single cell RNA-Seq data from 8 healthy lung samples were obtained from GSE128033 ^47^ and futher analyzed using R programming language. A total of 5898240 cells were filtered down to 21310 according to the following criteria: 1) number of genes per cell (nFeature, > 200 and < 6000); 2) Unique Molecular Identifiers (nCount, > 1000); and 3) % of mitochondrial genes (< 15 %). Data were then normalized using LogNormalize with a scale factor of 10000 and the top 2000 features were identified with the vst method using Seurat v.5.0.1. The resulting Seurat object was converted into an Anndata object using sceasy v.0.0.7 and the integration was processed using scVI v.1.0.4 and Scanpy v.1.9.8 in a Python enviroment. The trained model was re-converted into a Seurat object. The Seurat FindNeighbors function was used with the scVI reduction to obtain the resulting clusters with a resolution of 1.2. To focus on macrophages and monocytes, macrophage (*CD163*, *FABP4*, *LYVE1* as markers) and monocyte (*FCN1* as marker) clusters were identified and re-integrated using scVI v.1.0.4 in a Python enviroment and later re-clustered in Seurat v.5.0.1 with a resolution of 0.6.

### Phagocytosis assay

Phagocytosis ability was assessed by flow cytometry using pHrodo Red *E. coli* BioParticles Conjugates (Thermo Fisher), following the procedures recommended by the manufacturer. Macrophages were cultured in 24-well plate and exposed to pHrodo bioparticles for 60 min at 37 °C/5% CO2. Cells were then harvested and assessed by flow cytometry.

### Efferocytosis assay

Jurkat cells were cultured in RPMI 1640 medium without FCS for 16 h, and then treated with staurosporine (0.5 μg/ml, SIGMA), followed by an incubation at 37 °C for 3 h. Staurosporine treatment yielded a population of 83% annexin V⍰+⍰cells. Apoptotic cells (AC) were resuspended at a concentration of 1⍰×⍰10^6^ cells/ml and labeled with CellTrace™ Violet reagent (0.5 μM, Invitrogen, Thermo Scientific) for 20 min. For efferocytosis, macrophages were cultured with labeled AC (ratio 1:4) in p24 plates during 1 h at 37 °C. After 1 h, macrophages were rinsed with PBS to remove unbound AC and detached with PBS 5 mM EDTA, and pelleted by centrifugation before analyzing by flow cytometry.

### Bioenergetics profile

The XF24 extracellular flux analyzer (Seahorse Biosciences, North Billerica, MA) was used to determine the bioenergetic profile of intact cells. Briefly, cells were seeded (200,000 cells/well) in XF24 plates (Seahorse Biosciences) and allowed to recover for 24 h. Cells were then incubated in bicarbonate-free DMEM (Sigma-Aldrich) supplemented with 11.11 mM glucose, 2 mM L-glutamine, 1 mM pyruvate, and 2% FBS (Sigma-Aldrich) in a CO2-free incubator for 1 h. The oxygen consumption rate (OCR) and extracellular acidification rate (ECAR), a proxy for lactate production, were recorded to assess the mitochondrial respiratory activity and glycolytic activity, respectively. After four measurements under basal conditions, cells were treated sequentially with 1 μM oligomycin, 0.6 μM carbonyl cyanide p-(trifluoromethoxy)phenylhydrazone (FCCP), 0.4 μM FCCP, and 0.5 μM rotenone plus 0.5 mM antimycin A (Sigma-Aldrich), with three consecutive determinations under each condition that were subsequently averaged. Nonmitochondrial respiration (OCR value after rotenone plus antimycin A addition) was subtracted from all OCR measurements. ATP turnover was estimated from the difference between the basal and the oligomycin-inhibited respiration, and the maximal respiratory capacity was the rate in the presence of the uncoupler FCCP ^48^. Six independent replicas of each analysis were done, and results were normalized according to protein concentration.

### Multicolor Fluorescence Confocal Microscopy

Lung samples were collected from autopsies performed on patients who died from diagnosed SARS-CoV-2 infection or unrelated causes (myocardial infarction, as controls) at the Gregorio Marañón Hospital, Madrid. Patient data and samples were obtained in accordance with Ethics Committees of Instituto de Investigación Sanitaria Gregorio Marañón requirements. Formalin-fixed and paraffin-embedded tissues were deparaffinized, rehydrated, and unmasked by steaming in 10 mM sodium citrate buffer pH 9.0 (Dako Glostrup, Denmark) for 7 minutes. Slides were blocked with 5 μg/ml human immunoglobulins solved in blocking serum-free medium (Dako) for 30 minutes, and then sequentially incubated with 5–10 μg/ml primary antibodies, overnight at 4°C, and proper fluorescent secondary antibodies (Jackson Immunoresearch, West Grove, PA, US) for 1h. Washes were performed by immersion in PBS containing 0.05% Tween-20, and single-cell quantification was performed in 3–5 fields imaged with the ACS_APO 20x/NA 0.60 objective of a confocal microscope (Leica, SPE). Mean Fluorescence Intensity (MFI) of proteins of interest was obtained at segmented CD68+ macrophages using the ‘analyze particle’ plugging of ImageJ2 software ^49^, as previously shown ^50^.

### Statistical analysis

Statistical analyses were conducted using the GraphPad Prism software. For comparison of means, and unless otherwise indicated, statistical significance of the generated data was evaluated using the paired Student t test or One-Way ANOVA followed by multiple comparisons using the Dunnett’s test or Fisheŕs LSD test. In all cases, *p*<0.05 was considered as statistically significant.

## RESULTS

### Monocyte-derived macrophages generated in the presence of GM-CSF (GM-MØ) or M-CSF (M-MØ) differ in the state of activation of GSK3α and GSK3β

Monocyte-derived M-MØ exhibit a transcriptional profile akin to that of pathogenic macrophages in severe COVID-19 ^22^, whereas GM-MØ resemble lung alveolar macrophages (AMØ) ^22^. Our prior research has established that MAFB determines the transcriptional and functional specification of monocyte-derived M-MØ, characterized by significantly higher MAFB levels compared to GM-MØ ^27^. Since the protein levels of large MAF family members (MAF, MAFA, MAFB, NRL) are controlled through GSK3 phosphorylation-dependent proteasome degradation ^30^, we initially assessed the relative levels of the inhibitory (Ser^21^ for GSK3α and Ser^9^ for GSK3β) and activation-associated (Tyr^279^ for GSK3α and Tyr^216^ for GSK3β) phosphorylation of GSK3 in M-MØ and GM-MØ. Ser^21^-GSK3α and Ser^9^-GSK3β phosphorylation was more pronounced in M-MØ, aligning with their elevated levels of MAFB (Figure 1A). Conversely, no significant difference in Tyr^279^–GSK3α and Tyr^216^–GSK3β phosphorylation was observed between M-MØ and GM-MØ (Figure 1B). Therefore, the heightened expression of MAFB in M-MØ correlates with a higher inhibitory phosphorylation of GSK3.

**Figure 1.**
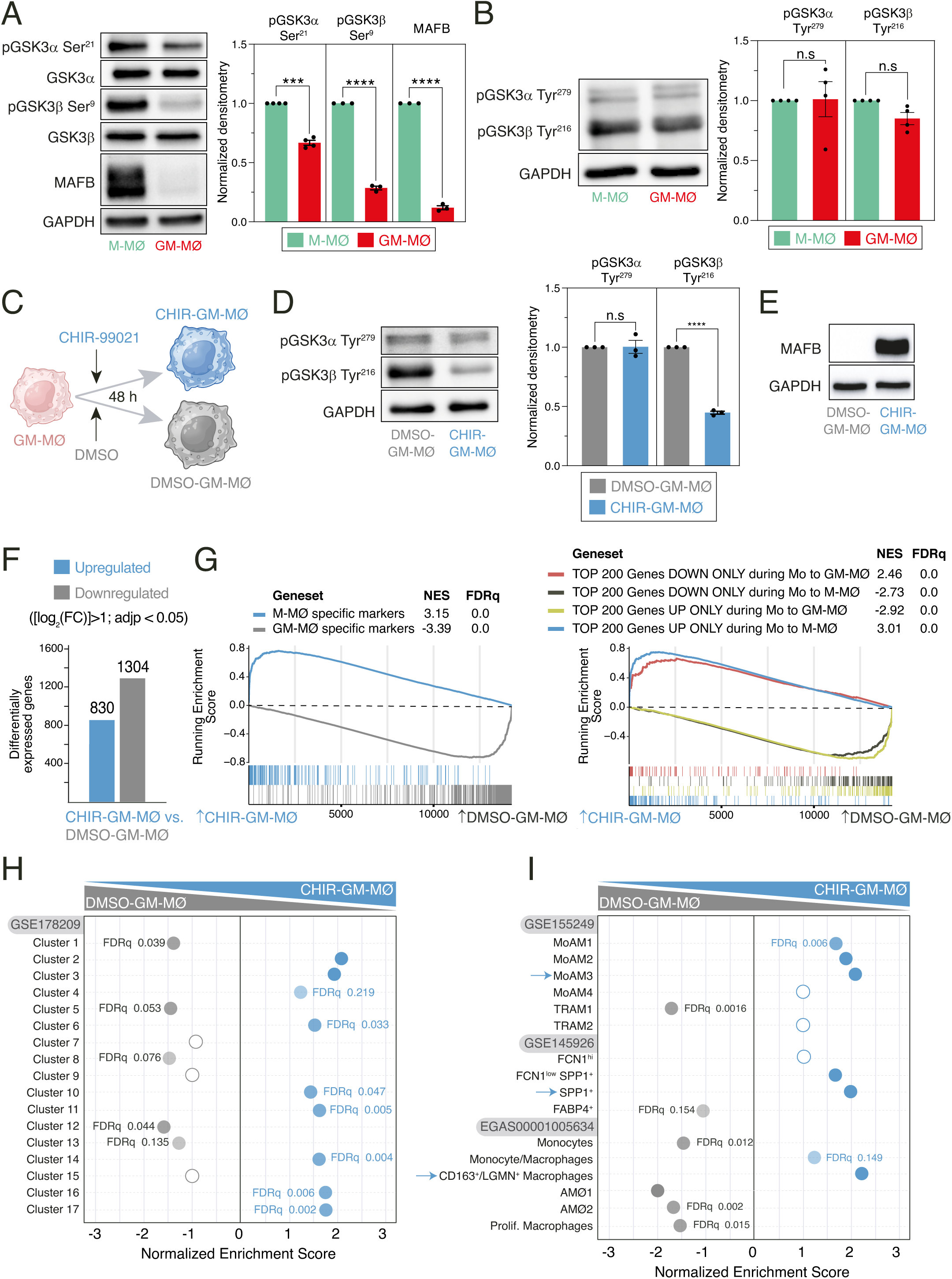
Transcriptional effect of GSK3 inhibition on monocyte-derived GM-MØ. **A**. MAFB, total GSK3α, total GSK3β, Ser^9^-phosphorylated GSK3β and Ser^21^-phosphorylated GSK3α levels in M-MØ and GM-MØ, as determined by Western blot (left panel). GAPDH protein levels were determined as protein loading control. Mean ± SEM of the MAFB/GAPDH, Ser^21^-phosphorylated GSK3α/total GSK3α and Ser^9^-phosphorylated GSK3β/total GSK3β protein ratios from three independent experiments are shown (right panel) (paired Student’s t test: ***, p<0.005; ****, p<0.001). A representative Western blot experiment is shown in each case in the upper panel. **B**. p-Tyr^279^– GSK3α and p-Tyr^216^–GSK3β levels in four independent samples of M-MØ and GM-MØ, as determined by Western blot (left panel). GAPDH protein levels were determined as protein loading control. Mean ± SEM of the p-Tyr^279^–GSK3α/GAPDH and p-Tyr^216^–GSK3β/GAPDH protein ratios from four independent experiments are shown (right panel) (paired Student’s t test: n.s., not significant). **C.** Schematic representation of the exposure of GM-MØ to CHIR-99021 for 48 hours. **D.** Tyr^279^–phosphorylated GSK3α and Tyr^216^–phosphorylated GSK3β levels in three independent preparations of DMSO- or CHIR-99021-treated GM-MØ, as determined by Western blot (left panel). GAPDH protein levels were determined as protein loading control. Mean ± SEM of the p-Tyr^216^–GSK3β/GAPDH and p-Tyr^279^–GSK3α/GAPDH protein ratios from the three independent experiments are shown (right panel) (paired Student’s t test: ****, p<0.001). **E**. MAFB protein levels in DMSO-GM-MØ and CHIR-GM-MØ, as determined by Western blot. A representative experiment is shown. **F**. Number of differentially expressed genes ([log2FC] >1; adjp<0.05) between DMSO-GM-MØ and CHIR-GM-MØ. Differential gene expression was assessed using DESeq2. **G**. GSEA of the indicated gene sets (from GSE68061, left panel; from GSE188278, right panel) on the ranked comparison of the CHIR-GM-MØ vs. DMSO-GM-MØ transcriptomes. Normalized Enrichment Score (NES) and FDRq values are indicated in each case. **H**. Summary of GSEA of the gene sets that define the tissue-resident monocyte and macrophage states (MoMac-VERSE) ^2^ on the ranked comparison of the CHIR-GM-MØ vs. DMSO-GM-MØ transcriptomes. FDRq values are indicated only if FDRq>0.0; empty dots, Not Significant. **I.** Summary of GSEA of the gene sets that characterize the macrophage subsets identified in severe COVID-19 ^23–25^ on the ranked comparison of the CHIR-GM-MØ vs. DMSO-GM-MØ transcriptomes. FDRq values are indicated only if FDRq>0.0; empty dots, Not Significant.

### Inhibition of GSK3 leads to transcriptional, phenotypic, functional, and metabolic reprogramming in monocyte-derived GM-MØ

Building upon the above findings, we aimed to examine the macrophage re-programming effects of GSK3 activity modulation in GM-MØ through the use of the GSK3 inhibitor CHIR-99021 (Figure 1C). Kinetics and dose-response analysis of the effects of CHIR-99021 on MAFB expression showed that maximal protein levels were achieved after a 24-48 hour exposure to 10μM CHIR-99021 (Supplementary Figure 1A), conditions that were used hereafter. Indeed, treatment of GM-MØ with CHIR-99021 for 48 hours resulted in a significant decrease in the activation-associated Tyr^216^–GSK3β phosphorylation (Figure 1D) and a robust elevation in MAFB protein levels (Figure 1E). Moreover, reversal of the Tyr^216^/Ser^9^ GSK3β phosphorylation ratio was also seen after CHIR-99021 treatment, albeit it was not observed after exposure to other GSK3 inhibitors (SB-216763 or LiCl; Supplementary Figure 1B). RNA-Seq analysis of CHIR-GM-MØ unveiled a huge transcriptional impact of GSK3 inhibition, with more than 2000 differentially expressed genes ([log_2_FC]>1; *adj p* < 0.05) (Figure 1F). Corresponding with the higher MAFB expression, Gene Set Enrichment Analysis (GSEA) showed that the CHIR-GM-MØ transcriptome was significantly enriched in genes that either mark M-MØ (GSE68061) or are upregulated during monocyte-to-M-MØ differentiation (Figure 1G). Conversely, GM-MØ-specific genes (GSE68061) or genes increased during monocyte-to-GM-MØ differentiation, were downregulated (Figure 1G). Additional gene ontology analysis further supported the pathophysiological significance of the macrophage re-programming induced by modulating GSK3 activity. Thus, analysis of the MoMac-VERSE (a resource that identified conserved monocyte and macrophage states derived from healthy and pathologic human tissues) (GSE178209) ^2^, indicated that GSK3 inhibition augments the expression of the gene sets that define MoMac-VERSE subsets identified as long-term resident macrophages [Cluster HES1_Mac (#2)] and tumor-associated macrophages with an M2-like signature [Clusters HES1_Mac (#2), TREM2_Mac (#3), C1Q^hi^_Mac (#16) and FTL_Mac (#17)] ^2^ (Figure 1H). Moreover, GSK3 inhibition increased the expression of the gene sets that define pathogenic pro-fibrotic macrophage subsets in severe COVID-19 [(Group 3 or SPP1^+^, GSE145926) ^24^, MoAM3 (GSE155249) ^25^ or CD163^+^/LGMN^+^ MØ (EGAS00001005634) ^23^] (Figure 1I). Therefore, GSK3 inhibition enhances MAFB expression and re-programs macrophage by promoting the acquisition of the gene profile that defines anti-inflammatory and pro-fibrotic macrophages.

In agreement with the increased MAFB levels detected in CHIR-GM-MØ (Figure 1E), screening for mediators of the re-programming action of GSK3 inhibition using DoRoThEA ^51^ (https://saezlab.github.io/dorothea/) revealed a significant enrichment in MAFB-regulated genes (Figure 2A), and similar results were yielded upon Enrichr analysis ^44,45^(https://maayanlab.cloud/Enrichr/) (Supplementary Figure 1C). In fact, GSEA confirmed that CHIR-GM-MØ are enriched in MAFB-dependent genes ^22,27^ (Figure 2B-C) along with macrophage genes whose regulatory regions are directly bound by MAFB ^52^ (75-geneset, Figure 2C), including *CCL2*, *CCL8*, *CCL18*, *IGF1*, *IL10* and *LGMN* (Figure 2D). Importantly, all these changes were reflected in the phenotype of CHIR-GM-MØ, that showed elevated expression of transcription factors like MAF (Figure 2E), cell surface proteins like CD163 and FOLR2 (Figure 2E, F), and soluble factors like CCL2, CCL18, IL-10, LGMN and SPP1 (Figure 2G), all of which are either direct MAFB targets ^52^ or MAFB-regulated genes ^22,27^. Likewise, the attenuated expression of the GM-MØ-specific *INHBA* gene (Figure 2D) corresponded to a decrease in Activin A production by CHIR-GM-MØ (Figure 2G). These results underscore the phenotypic re-programming induced by GSK3 inhibition in GM-MØ, wherein there is an upregulation of MAFB-dependent transcription factors, cell surface markers, and soluble factors which collectively define an anti-inflammatory/pro-fibrotic phenotype characteristic of M-CSF-dependent monocyte-derived macrophages.

**Figure 2.**
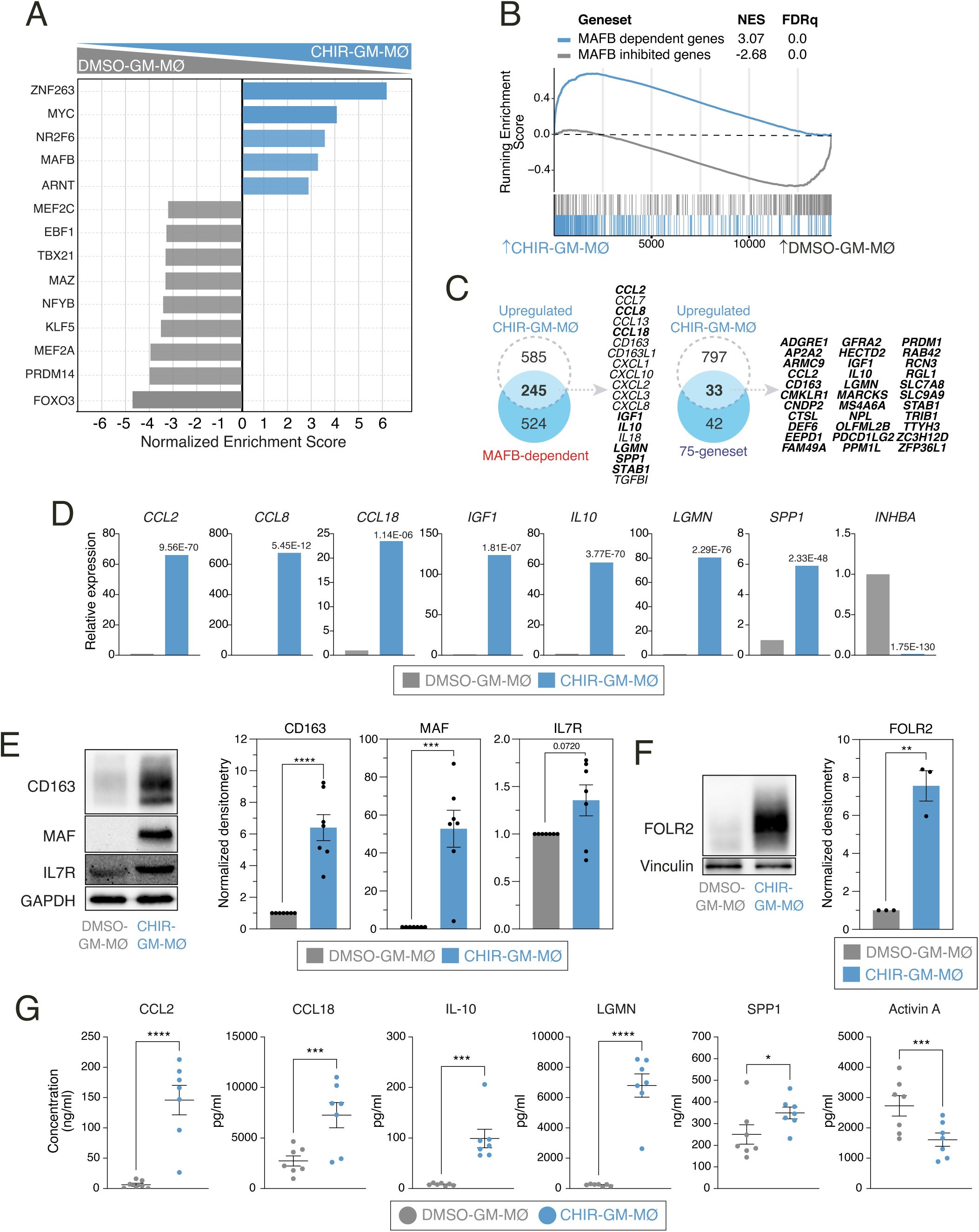
Phenotypic effects of GSK3 inhibition on monocyte-derived GM-MØ: Enhanced expression of MAFB-dependent genes and proteins. **A.** DoRoThEA ^51^ analysis on the ranked comparison of the DMSO-GM-MØ and CHIR-GM-MØ transcriptomes. **B**. GSEA of MAFB-regulated gene sets (from GSE155719) on the ranked comparison of the DMSO-GM-MØ and CHIR-GM-MØ transcriptomes. NES and FDRq values are indicated in each case **C**. Overlap between the genes upregulated (|log2FC|>1; adjp<0.05) in CHIR-GM-MØ (relative to DMSO M-MØ) and MAFB-dependent genes (from GSE155719, left panel) or the 75-geneset of MAFB-regulated genes (right panel) ^52^, with indication of some of the overlapping genes. **D**. Relative mRNA levels of the indicated genes in DMSO-GM-MØ and CHIR-GM-MØ, as determined by RNA-Seq on three independent samples (GSE256208). *Adjp* of the comparison, shown in each case, was assessed using DESeq2. **E**. CD163, MAF and IL7R protein levels in DMSO-GM-MØ and CHIR-GM-MØ, as determined by Western blot (left panel). GAPDH protein levels were determined as protein loading control. Mean ± SEM of the MAF/GAPDH, IL7R/GAPDH and CD163/GAPDH protein ratios from seven independent experiments are shown (right panels) (paired Student’s t test: ***, p<0.005; ****, p<0.001). A representative Western blot experiment is shown in each case. **F**. FOLR2 protein levels in DMSO-GM-MØ and CHIR-GM-MØ, as determined by Western blot (left panel). Vinculin protein levels were determined as protein loading control. Mean ± SEM of the FOLR2/Vinculin protein ratio from three independent experiments are shown (lower panel) (paired Student’s t test: **, p<0.01). A representative Western blot experiment is shown. **G**. Production of the indicated soluble factors by DMSO-GM-MØ and CHIR-GM-MØ, as determined by ELISA. Mean ± SEM of seven independent samples are shown (paired Student’s t test: *, p<0.05; **, p<0.01; ***, p<0.005; ****, p<0.001).

Remarkably, the phenotypic changes induced by GSK3 inhibition were also accompanied by alterations in macrophage functional capabilities. Thus, CHIR-GM-MØ exhibited elevated anti-inflammatory activity, evidenced by increased production of IL-10 upon activation by inflammatory cytokines such as TNF or IFNγ, or exposure to PAMPs like LPS or the TLR7 ligand CL264 (Figure 3A). In addition, CHIR-GM-MØ demonstrated increased phagocytic ability for *E. coli* particles (Figure 3B), as well as enhanced efferocytosis (Figure 3C), with both activities being typically higher in M-MØ. Collectively, these findings indicate that GSK3 inhibition re-educates GM-MØ macrophages by facilitating the acquisition of transcriptional, phenotypic, and functional properties characteristic of M-CSF-dependent monocyte-derived macrophages. Of note, in line with the well-established connection between inflammatory potential and metabolic state in macrophages ^53^, GSK3 inhibition also induced modifications in various metabolic parameters in GM-MØ, including basal respiration and maximal ATP production (Supplementary Figure 1D-F).

**Figure 3.**
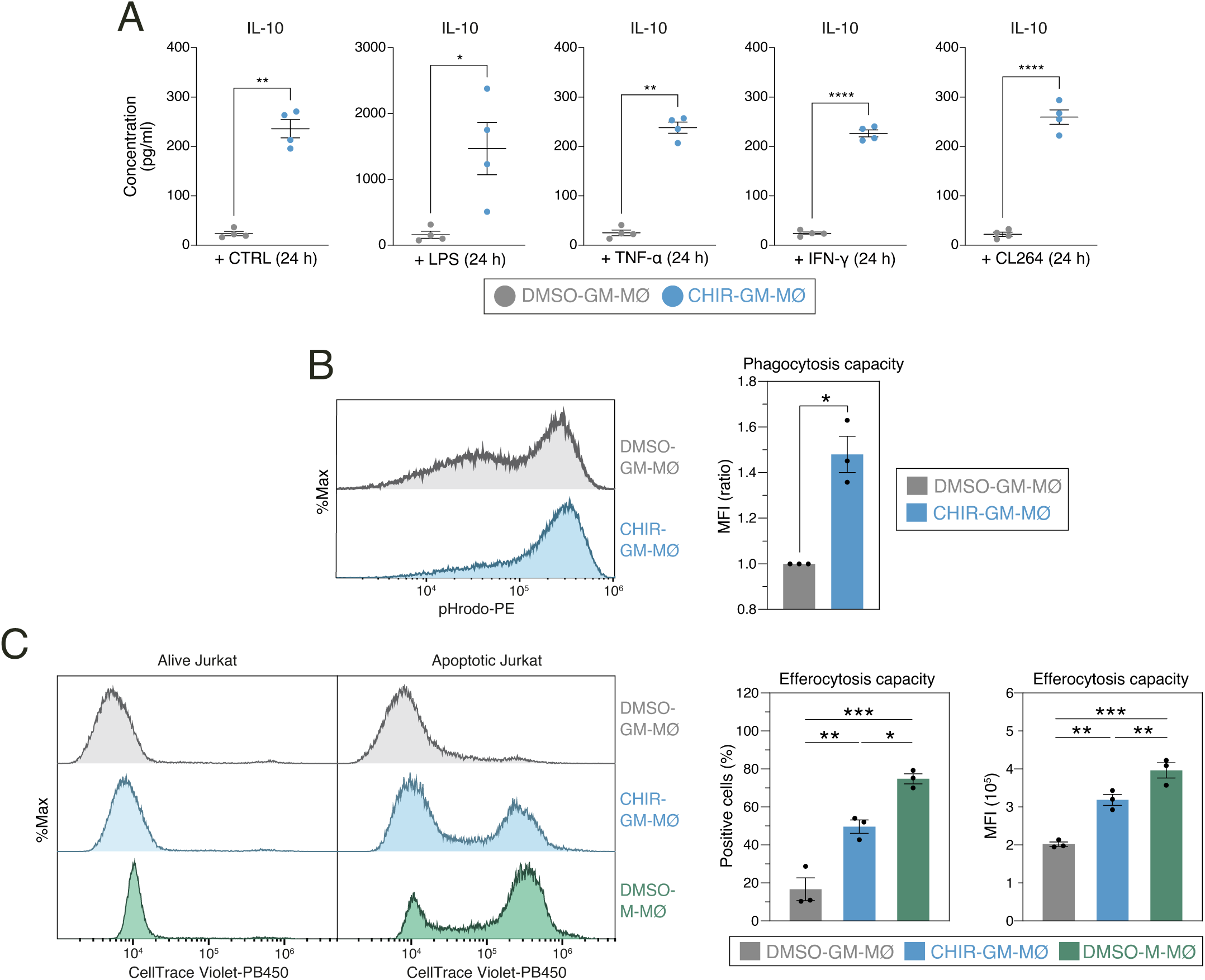
Functional consequences of GSK3 inhibition on monocyte-derived GM-MØ. **A.** Production of IL-10 by untreated (−) or stimulated (LPS, TNF, IFNγ or CL264) DMSO-GM-MØ and CHIR-GM-MØ, as determined by ELISA. Mean ± SEM of four independent samples are shown (paired Student’s t test: *, p<0.05; **, p<0.01; ***, p<0.005; ****, p<0.001). **B.** Phagocytosis of pHRodo-labeled bacterial particles by DMSO-GM-MØ and CHIR-GM-MØ, as determined by flow cytometry. Mean ± SEM of three independent samples are shown (paired Student’s t test: *, p<0.05). A representative flow cytometry analysis is shown in the left panel. **C**. Efferocytosis capacity of DMSO-GM-MØ, CHIR-GM-MØ and DMSO-M-MØ, as determined by flow cytometry using staurosporine-induced CellTrace Violet-labeled apoptotic Jurkat cells. The percentage of positive cells and mean fluorescence intensity are shown. Mean⍰±⍰SEM of 3 independent samples are shown (one-way ANOVA with Fisher LSD test: *, p<0.05; **, p⍰<⍰0.01; ***, p⍰<⍰0.005). Representative flow cytometry histograms for CellTrace Violet emission of DMSO-GM-MØ, CHIR-GM-MØ and DMSO-M-MØ are shown.

### siRNA-mediated knock-down of *GSK3A* and/or *GSK3B* enhances the expression of the MAFB-dependent profiles of M-MØ and pathogenic pro-fibrotic lung macrophages

To fully support the involvement of GSK3 in the macrophage re-programming action of CHIR-99021, we next assessed the transcriptional consequences of knocking-down *GSK3A* (coding for GSK3α) and/or *GSK3B* (coding for GSK3β) in GM-MØ (Figure 4A,B). Like CHIR-99021, silencing of both *GSK3A* and *GSK3B* augmented the expression of MAFB, with the simultaneous silencing of both *GSK3A* and *GSK3B* genes having a stronger effect (Figure 4B), and modulated the expression of 329 genes (Figure 4C,D). Of note, silencing of *GSK3A* and *GSK3B* reproduced the effects of CHIR-99021, as it strongly enhanced the expression of genes upregulated by the inhibitor in either GM-MØ or *ex vivo* isolated Alveolar Macrophages (Figure 4E and not shown). *GSK3A/B* knock-down triggered a significant enrichment in the expression of MAFB-dependent genes (Figure 4F), genes acquired along the monocyte-to-M-MØ differentiation (Figure 4G), and genes that mark M-MØ, while reducing the expression of GM-MØ-specific genes (Figure 4G). Furthermore, in agreement with the effect of CHIR-99021, *GSK3A/B* knock-down enhanced the expression of the MoMac-VERSE macrophage clusters HES1_Mac (#2) and TREM2_Mac (#3) ^2^ (Figure 4H) as well as the gene sets that define pathogenic pro-fibrotic macrophage subsets in severe COVID-19 [(SPP1^+^, GSE145926)^24^, MoAM3 (GSE155249) ^25^ or CD163^+^/LGMN^+^ MØ (EGAS00001005634) ^23^] (Figure 4I, Supplementary Figure 2A). Conversely, silencing of *GSK3A* and *GSK3B* drastically reduced the expression of the gene sets that characterize tissue-resident lung Alveolar Macrophages in COVID-19, with the concomitant silencing of *GSK3A* and *GSK3B* again showing a stronger effect (Figure 4I). Altogether, these results support the relevance of GSK3α and GSK3β as relevant targets for human macrophage re-programming.

**Figure 4.**
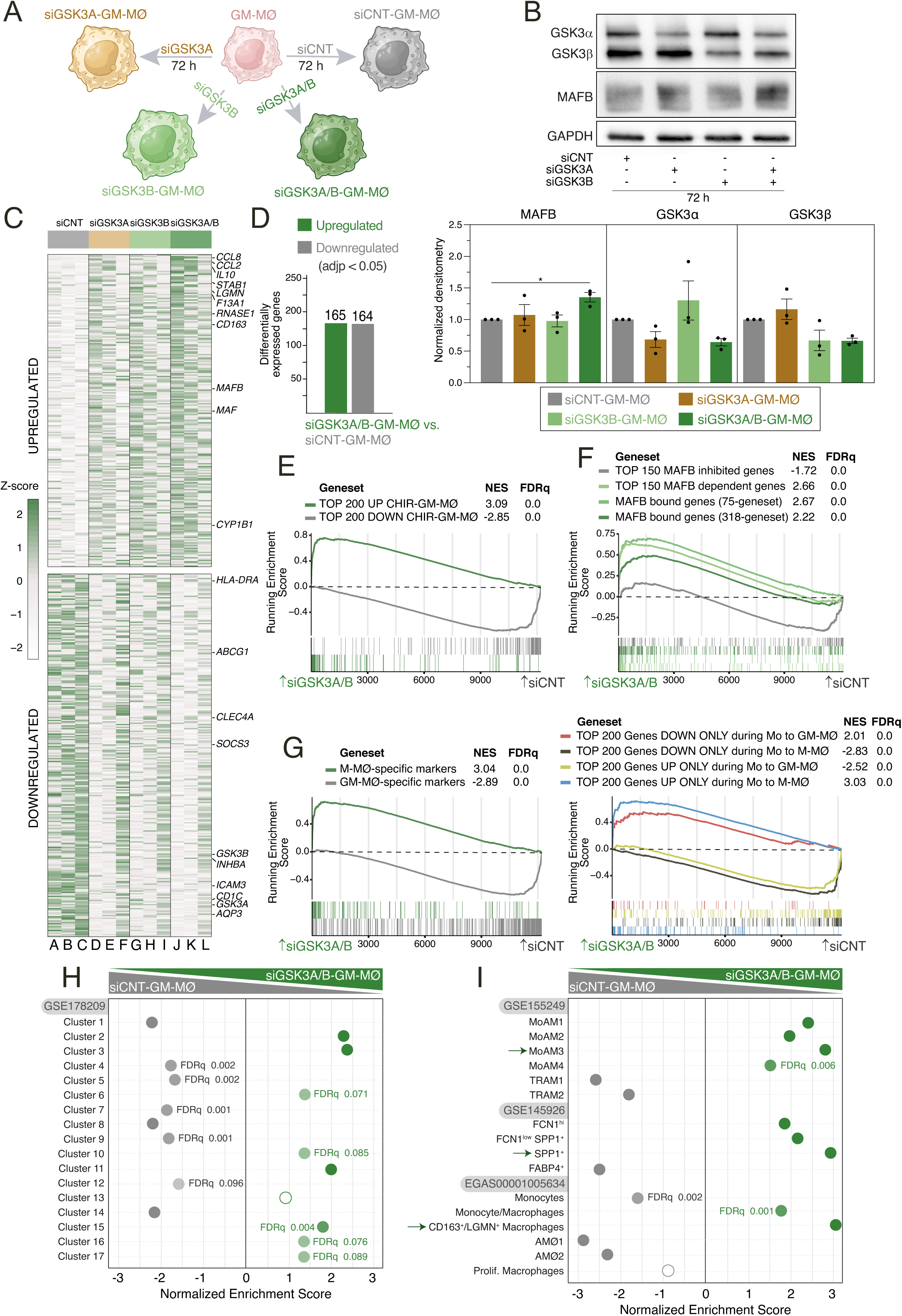
Transcriptional effects of GSK3 knock-down in monocyte-derived GM-MØ. **A.** Schematic representation of the siRNA-mediated GSK3 knock-down procedure in monocyte-derived GM-MØ. **B**. Total GSK3α, GSK3β and MAFB protein levels in monocyte-derived GM-MØ after siRNA-mediated *GSK3A* and/or *GSK3B* silencing, as determined by Western blot (upper panel). GAPDH protein levels were determined as protein loading control. Mean ± SEM of GSK3β/GAPDH and GSK3α/GAPDH protein ratios from the three independent experiments are shown (lower panel) (one-way ANOVA with Dunnet test:*, p<0.05). **C**. Heatmap of the relative expression of the genes significantly altered after *GSK3A* and *GSK3B* knock-down in siCNT-GM-MØ (lanes A-C), siGSK3A-GM-MØ (lanes D-F), siGSK3B-GM-MØ (lanes G-I), siGSK3A/B-GM-MØ (lanes J-L). Representative genes are indicated. **D**. Number of differentially expressed genes (adjp<0.05) between siGSK3A-GM-MØ, siGSK3B-GM-MØ or siGSK3A/B-GM-MØ and siCNT-GM-MØ. Differential gene expression was assessed using DESeq2. **E**. GSEA of the genes whose expression is significantly modulated by CHIR-99021 in either GM-MØ (from Figure 1, this manuscript) or in *ex vivo* isolated Alveolar Macrophages (AMØ, see below in Figure 6) on the ranked comparison of the siGSK3A/B-GM-MØ vs. siCNT-GM-MØ transcriptomes. **F.** GSEA of the indicated gene sets (from GSE68061) on the ranked comparison of the siGSK3A/B-GM-MØ vs. siCNT-GM-MØ transcriptomes. **G**. Summary of GSEA of the indicated gene sets (from GSE188278) on the ranked comparison of siGSK3A/B-GM-MØ vs. siCNT-GM-MØ. **H.** Summary of GSEA of the gene sets that define the tissue-resident monocyte and macrophage states (MoMac-VERSE) ^2^ on the ranked comparison of the siGSK3A/B-GM-MØ vs. siCNT-GM-MØ transcriptomes. FDRq values are indicated only if FDRq>0.0; empty dots, Not Significant. **I.** Summary of GSEA of the gene sets that characterize the macrophage subsets identified in severe COVID-19 ^23–25^ on the ranked comparison of siGSK3A/B-GM-MØ vs. siCNT-GM-MØ. FDRq values are indicated only if FDRq>0.0; empty dots, Not Significant.

### GSK3 inhibition redirects monocytes towards acquiring a gene profile reminiscent of anti-inflammatory M-MØ

Given the swift and profound anti-inflammatory impact observed in fully differentiated GM-MØ upon GSK3 inhibition or knock-down, we next delved into assessing whether modulation of GSK3 activity influence the differentiation capability of peripheral blood monocytes. To that end, monocytes were exposed to CHIR-99021, and the resulting cells (CHIR-Mon, Figure 5A) exhibited increased MAFB levels as early as 16 hours post-treatment (Figure 5B). Remarkably, the 16-hour exposure to the GSK3 inhibitor induced significant alterations in the expression of over 2000 genes ([log_2_FC]>1; *adj p* < 0.05) (Figure 5C) and sufficed to augment the expression of M-MØ-specific genes and genes upregulated during M-CSF-dependent monocyte-to-M-MØ differentiation (Figure 5D), including *CCL2*, *CD163* and *IL10* (Figure 5E). Notably, the transcriptome of CHIR-Mon exhibited a significant enrichment in MAFB-dependent genes (Figure 5F, Supplementary Figure 2B) and genes characterizing the MAFB+ pathogenic macrophages in severe COVID-19 (Supplementary Figure 2C), whereas it showed an under-representation of genes inhibited by MAFB ^52^ (Figure 5F). Compared to DMSO, CHIR-99021 treatment upregulated the expression of 25% of M-MØ-specific genes, including MAFB-dependent genes like *CCL2*, *IL10* and *CD163*, while downregulating 30% of GM-MØ-specific genes (Figure 5G), and this transcriptional effect was also evident in comparison with M-CSF-treated monocytes (Supplementary Figure 2D). To broaden the scope of these findings, we also examined the consequences of inhibiting GSK3 in monocytes exposed to a pathologic pro-inflammatory environment, namely synovial fluid from patients with active rheumatoid arthritis (RASF) (Figure 5H). Evaluation of CHIR-99021-treated monocytes in the presence of RASF revealed an enhanced expression of MAFB (Figure 5I) and MAFB-dependent genes such as *IL10* and *LGMN* (Figure 5J). Therefore, GSK3 inhibition in monocytes also leads to heightened MAFB expression, ultimately steering differentiation towards anti-inflammatory monocyte-derived macrophages.

**Figure 5.**
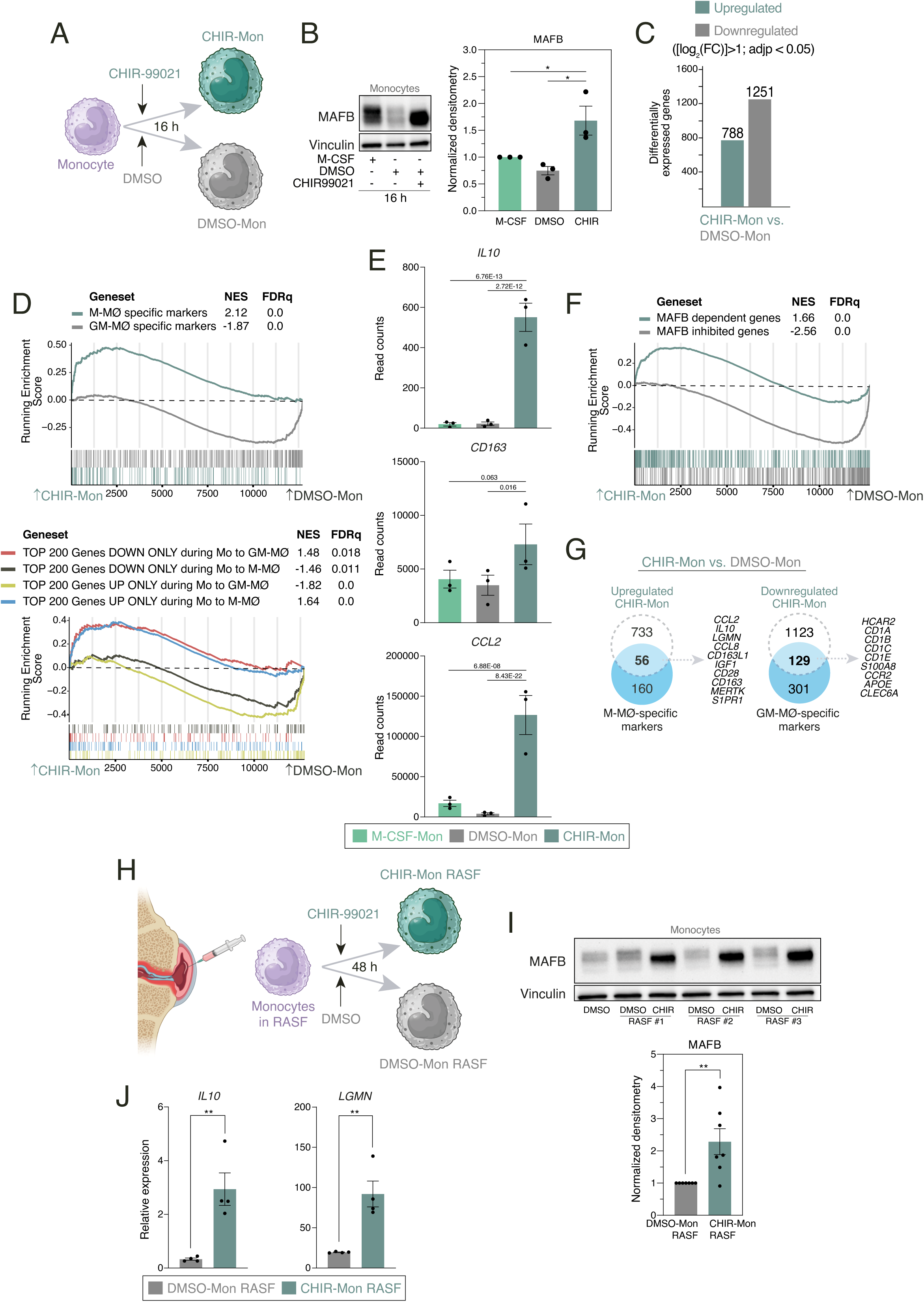
Transcriptional effects of GSK3 inhibition on human peripheral blood monocytes. **A.** Schematic representation of the exposure of human monocytes to CHIR-99021 or DMSO for 16h. **B**. MAFB protein levels in monocytes exposed for 16h to DMSO, CHIR-99021 or M-CSF, as determined by Western blot (left panel). Vinculin protein levels were determined as protein loading control. Mean ± SEM of the MAFB/Vinculin protein ratio from the three independent experiments shown (right panel) (one-way ANOVA with Fisher LSD test: ***, p<0.005). **C**. Number of differentially expressed genes ([log2FC] >1; adjp<0.05) between DMSO-Mon and CHIR-Mon. Differential gene expression was assessed using DESeq2. **D**. GSEA of the indicated gene sets (from GSE68061, upper panel; from GSE188278, lower panel) on the ranked comparison of the DMSO-Mon and CHIR-Mon transcriptomes. NES and FDRq values are indicated in each case. **E**. Relative mRNA levels of the indicated genes in monocytes exposed for 16h to DMSO, CHIR-99021 or M-CSF, as determined by RNA-Seq on three independent samples (GSE256538). *Adjp* of the comparison, shown in each case, was assessed using DESeq2. **F**. GSEA of MAFB-regulated gene sets (from GSE155719) on the ranked comparison of the DMSO-Mon and CHIR-Mon transcriptomes. NES and FDRq values are indicated in each case. **G**. Overlap between the genes upregulated (|log2FC|>1; adjp<0.05) in CHIR-Mon (relative to DMSO-Mon) and M-MØ-specific marker genes (from GSE68061) (left panel) and genes downregulated (|log2FC|>1; adjp<0.05) in CHIR-Mon (relative to DMSO-Mon) and GM-MØ-specific marker genes (from GSE68061) (right panel), with indication of some of the overlapping genes. **H**. Schematic representation of the exposure of human monocytes to synovial fluid from rheumatoid arthritis patients (RASF) for 48h in the presence or absence of CHIR-99021. **I**. MAFB protein levels in monocytes exposed to three independent samples of RASF in the presence of DMSO or CHIR-99021, as determined by Western blot (upper panel). Vinculin protein levels were determined as protein loading control. Mean ± SEM of the MAFB/Vinculin protein ratio from seven independent experiments using three monocyte preparations and three unrelated RASF (lower panel) (paired Student’s t test: **, p<0.01). **J**. Relative expression of *IL10* and *LGMN* in monocytes exposed to RASF in the presence of DMSO or CHIR-99021, as determined by RT-PCR. Mean ± SEM of the results from the three independent samples are shown (paired Student’s t test: **, p<0.01).

### Inhibition of GSK3 in *ex vivo* isolated human alveolar macrophages promotes the acquisition of an M-MØ-like anti-inflammatory profile

To further validate the macrophage re-programming potential of GSK3 inhibition, we next conducted experiments on *ex vivo* isolated human alveolar macrophages (AMØ, Figure 6A), whose development and gene expression profile is GM-CSF-dependent ^17–19^. To that end, we determined the transcriptome of three independent isolates of AMØ, purified from broncho-alveolar lavages, after exposure to CHIR-99021 for 24 hours (Figure 6A). A total of 865 differentially expressed genes ([log_2_FC]>1; *adj p* < 0.05) were identified, with 316 genes upregulated in CHIR-AMØ (Figure 6B). Further comparisons and gene ontology analysis revealed that GSK3 inhibition by CHIR-99021 (CHIR-AMØ) upregulated the gene sets that define pathogenic pro-fibrotic macrophage subsets in severe COVID-19 [(SPP1^+^, GSE145926) ^24^, MoAM3 (GSE155249) ^25^ or CD163^+^/LGMN^+^ MØ (EGAS00001005634) ^23^] (Figure 6C,D; Supplementary Figure 2E). Conversely, CHIR-AMØ exhibited a reduced expression of the gene sets that characterize lung-resident AMØ in severe COVID-19 [AMØ1+AMØ2 clusters from EGAS00001005634) ^23^, FABP4+ from GSE145926) ^24^, TRAM1 from GSE155249 ^25^] (Figure 6D,E; Supplementary Figure 2F), including key AMØ-specific genes like *FABP4* and *MARCO* (Figure 6E), and similar resuts were seen upon analysis of AMØ-specific gene sets from other studies ^25,54^ (Supplementary Figure 2F). Like in the case of CHIR-GM-MØ, CHIR-AMØ exhibited an over-representation of M-MØ-specific genes and genes acquired along the monocyte-to-M-MØ differentiation (GSE188278) (Supplementary Figure 2G). Consistent with these findings, CHIR-AMØ exhibited higher expression of MAFB (Figure 6F) whose increase correlated with an augmented secretion of Legumain, CCL2 and IL-10 (Figure 6G), although the latter was only seen in two samples, probably reflecting heterogeneity in primary cell responses. In contrast, lower levels of activin A were detected in CHIR-AMØ-conditioned medium (Figure 6G). Remarkably, and in agreement with the shift in macrophage subsets that takes place in severe COVID-19 lungs (loss of AMØ, appearance of monocyte-derived pro-fibrotic macrophages) ^22–25^, the level of phosphorylated Ser^21^-GSK3α and Ser^9^-GSK3β was very significantly augmented in lung macrophages (CD68+) from severe COVID-19 patients, an increase that correlated with a robust enhancement of the macrophage expression of both MAFB and the MAFB-dependent protein CD163 (Figure 6H). These results collectively demonstrate that macrophage re-programming through GSK3 inhibition can also be achieved on *in vivo* isolated macrophages, leading to enhanced expression of MAFB and MAFB-dependent genes, the acquisition of an anti-inflammatory transcriptional profile and the loss of the signature of lung-resident human AMØ.

**Figure 6.**
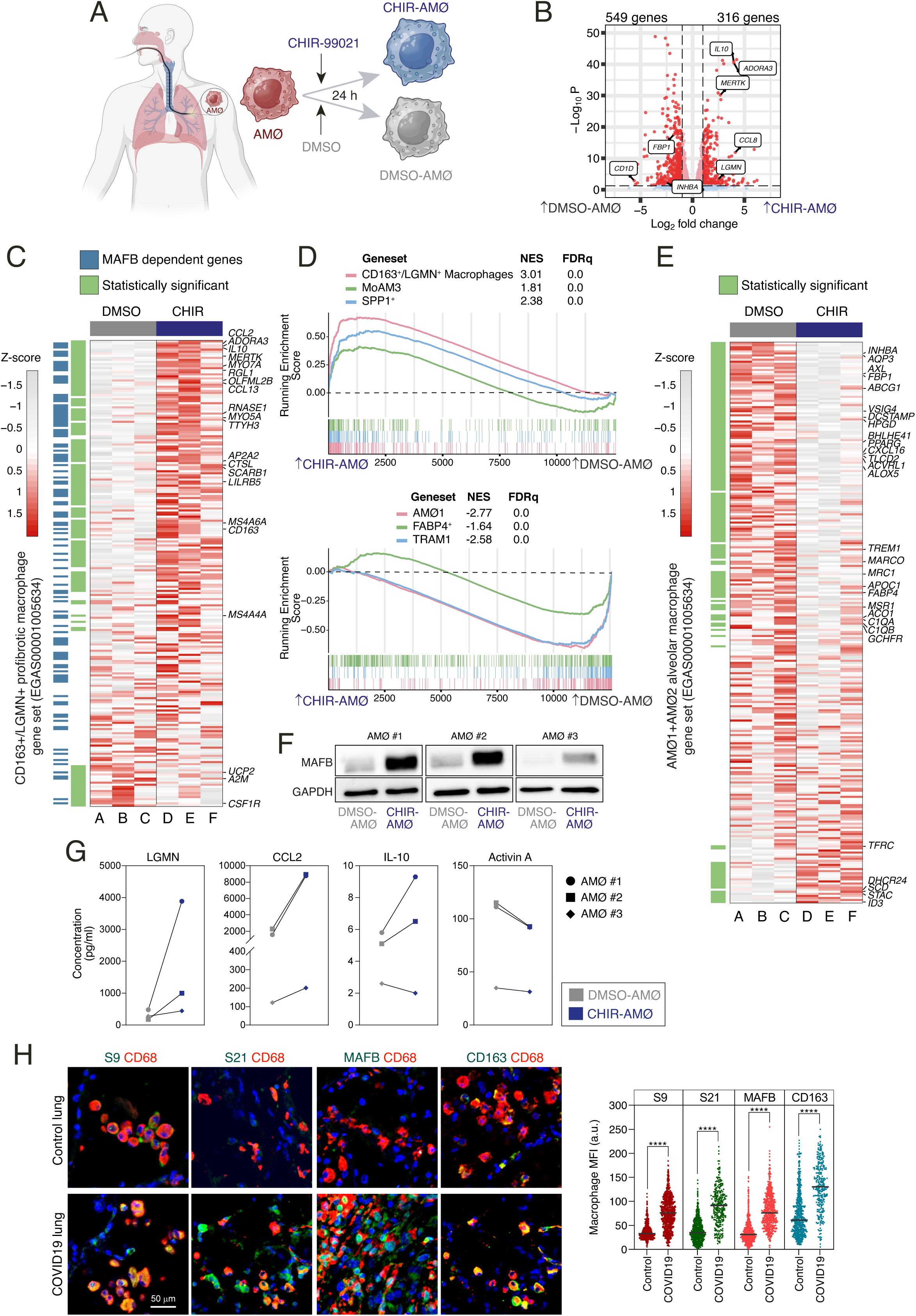
Transcriptional and phenotypic effects of GSK3 inhibition on *ex vivo* isolated human alveolar macrophages. **A**. Schematic representation of the exposure of *ex vivo* isolated human alveolar macrophages to CHIR-99021 or DMSO for 24h. **B**. Volcano plot depicting the differentially expressed genes between CHIR-AMØ and DMSO-AMØ. **C**. Heatmap of the relative expression of the genes within the CD163+/LGMN+ pro-fibrotic monocyte-derived macrophage cluster (from EGAS00001005634) ^23^ in DMSO-AMØ (lanes A-C) and CHIR-AMØ (lanes D-F). Differentially expressed between CHIR-AMØ and DMSO-AMØ are shown in greeen, and MAFB-dependent genes (GSE155719) are shown in blue. Representative genes are indicated. **D**. (Upper panel) GSEA of the gene sets that characterize the monocyte-derived profibrotic macrophage subsets identified in severe COVID-19, namely CD163+/LGMN+ (EGAS00001005634) ^23^, MoAM3 (GSE155249) ^25^ and SPP1+ (GSE145926) ^24^, on the ranked comparison of the transcriptomes of CHIR-AMØ vs. DMSO-AMØ. NES and FDRq values are indicated in each case. (Lower panel) GSEA of the gene sets that characterize tissue-resident alveolar macrophages in severe COVID-19, namely AMØ1 (EGAS00001005634) ^23^, TRAM1+ (GSE155249) ^25^ and FABP4+ (GSE145926) ^24^, on the ranked comparison of the transcriptomes of CHIR-AMØ vs. DMSO-AMØ. **E.** Heatmap of the relative expression of the genes within the Alveolar Macrophage signature (including all the genes overexpressed in the AMØ1 and AMØ2 macrophages clusters from EGAS00001005634) ^23^ in DMSO-AMØ (lanes A-C) and CHIR-AMØ (lanes D-F). Differentially expressed between CHIR-AMØ and DMSO-AMØ are indicated in greeen, and selected genes are shown. **F**. MAFB protein levels in three independent samples of *ex vivo* isolated AMØ exposed to DMSO or CHIR-99021 for 24h, as determined by Western blot. GAPDH protein levels were determined as protein loading control. **G**. Production of the indicated soluble factors by three independent preparations of DMSO-AMØ and CHIR-AMØ, as determined by ELISA. **H**. Expression of inactive GSK3 (S9, pSer^9^-GSK3β; S21, pSer^21^-GSK3α), MAFB and CD163 in human lung macrophages. Representative human lung tissues from COVID (n= 2) and control (n= 3) patients co-stained for CD68 (macrophage marker, red), Ser^21^-phosphorylated GSK3α, Ser^9^-phosphorylated GSK3β, MAFB or CD163 (green), and DAPI (nuclei, blue), as indicated (Scale bar, 50 μm) (left panel). Plots show MFI (in arbitrary units, a.u.) of single-cell CD68+ macrophages stained for each antibody (right panel) (paired Student’s t test: ****, p<0.001).

### GSK3- and MAFB-dependent genes are differentially expressed between human lung alveolar and interstitial macrophages

To assess whether GSK3 inhibition also modulates the phenotypic differences between human lung macrophage subsets in a physiological setting, we next took advantage of the available information on human healthy lungs obtained by single cell-RNA sequencing ^47^. Re-analysis of the single cell-RNA sequencing data ^47^ (Figure 7A; Supplementary Figure 3A-D) revealed several clusters expressing macrophage-associated markers (*CD163*, *FABP4*, *LYVE1, FCN1*) (Supplementary Figure 3D), and whose subsequent re-clustering identified cell subsets expressing previously defined markers for infiltrating monocytes (Cluster 4/7/12:*FCN1*) ^23^, Alveolar macrophages (AMØ) (Clusters 0/2/5/13: *FABP4 TYMS*-negative; Cluster 9*: MARCO*, *INHBA; Cluster 11: PPARG*), proliferating AMØ (Cluster 15: *TYMS, MKI67, TOP2A, NUSAP1*) ^23^, and Interstitial Macrophages (IMØ) that variably express markers for tissue-resident macrophages like *LYVE1, RNASE1* and *LGMN* (Clusters 1/3/6/8/10/14) ^1,23,47^ (Figure 7B; Supplementary Figure 4A-C). The comparison of IMØ and AMØ subsets revealed that the genes upregulated upon inhibition of GSK3 include numerous MAFB-dependent genes (e.g., *MAFB*, *IL10*, *LGMN*, *SPP1*, *CCL2*) and showed a higher expression in IMØ (Figure 7C). On the other hand, the genes downregulated by CHIR-99021 were preferentially or exclusively expressed in AMØ, including *INHBA*, *GSN*, *FBP1*, *PPARG*, *AXL* and *AQP3* (Figure 7C). Therefore, GSK3 inhibition prompts macrophages towards the acquisition of a gene profile that resembles that of IMØ, while provokes the loss of the AMØ gene signature. Such a conclusion was also reached upon analysis of pseudo bulk-RNA sequencing data from GSE128033 ^47^. Indeed, numerous MAFB-dependent and GSK3-regulated genes were found within the 712 genes that are differentially expressed ([log_2_FC]>1; *adj p* < 0.05) between IMØ and AMØ (Figure 7D). Further GSEA evidenced that GSK3 inhibition in either GM-MØ (CHIR-GM-MØ) (Figure 7E) or AMØ (CHIR-AMØ) (Figure 7F) very significantly enhance the expression of MAFB-dependent IMØ-specific genes (as defined by Morse *et al*. ^47^ or by Li *et al*. ^55^), while simultaneosuly impaired the expression of genes characterizing human AMØ (Figure 7E-F). More importantly, a similar effect was observed after *GSK3A/B* silencing in GM-MØ, that yielded an enhanced expression of IMØ-specific genes and a downregulation of the gene set that defines AMØ (Figure 7G). Altogether, these results indicate that inhibiting GSK3 shifts the gene profile of Alveolar Macrophages towards that of Interstitial Macrophages (or monocyte-derived macrophages), and imply that the activation status of GSK3 significantly influences the specification of lung-resident macrophages (AMØ *vs*. IMØ).

**Figure 7.**
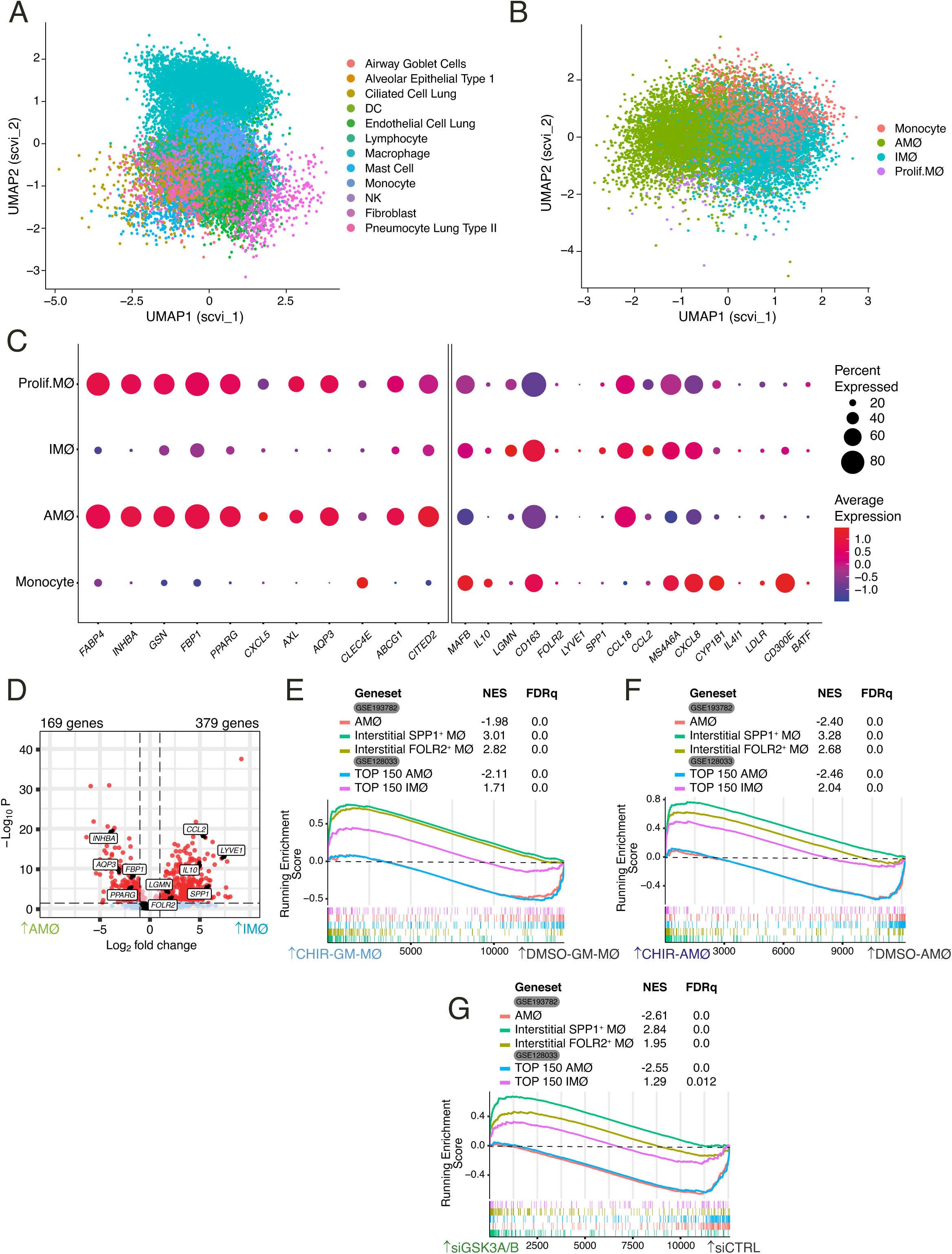
Differential expression of GSK3-regulated genes in human lung interstitial and alveolar macrophages. **A**. Embedding of 21310 single-cell transcriptomes from human lungs in UMAP (Uniform Manifold Approximation and Projection) with the scVI reduction. Cell-type annotation is based on expression of canonical marker genes, as reported in ^47^. **B**. Two-dimensional embedding computed by UMAP with the scVI reduction on computationally identified macrophages after filtering according to number of genes per cell (nFeature, > 200 and < 6000), Unique Molecular Identifiers (nCount, > 1000) and % of mitochondrial genes (< 15 %). **C**. Relative expression of the indicated genes in the four macrophage subsets defined upon re-clustering of the macrophages identified in the single cell RNA sequencing reported in GSE128033 ^47^. **D**. Volcano plot illustrating the differentially expressed genes between the Interstitial Macrophage (IMØ) and Alveolar Macrophage (AMØ) subsets defined upon re-analysis of the single cell RNA sequencing reported in GSE128033 ^47^. **E-G**. GSEA of the gene sets that define human lung AMØ or IMØ macrophage subsets (GSE128033 and GSE193782) ^47,55^ on the ranked comparison of CHIR-GM-MØ vs. DMSO-GM-MØ transcriptomes (**E**), CHIR-AMØ vs. DMSO-AMØ transcriptomes (**F**) or siGSK3A/B-GM-MØ vs. siCNT-GM-MØ (**G**).

## DISCUSSION

Our previous investigations have demonstrated that the macrophage re-programming effect of LXR modulators ^38,56^ and methotrexate ^37^ coincide with alterations in the protein levels of MAFB. Similarly, intravenous immunoglobulins have been shown to alter the inflammatory state of macrophages and modulate MAFB levels ^57^. Building upon these findings and others, we have explored whether modulating the activity of GSK3, a primary regulator of MAFB protein levels and activity, is sufficient to induce a shift in the inflammatory profile of human macrophages. Our results indicate that GSK3 inhibition impairs the expression of genes associated with GM-CSF-dependent differentiation of monocyte-derived macrophages while concurrently promoting the acquisition of genes and functions associated with M-CSF-dependent monocyte-derived macrophages, particularly those regulated by MAFB. The physiological relevance of these findings is underscored by the contrasting effect that GSK3 inhibition has on genes that define lung-resident alveolar macrophages (decrease) versus monocyte-derived macrophages recruited into the lung (increase) during inflammatory responses. Thus, modulation of GSK3 activity emerges as a plausible strategy for macrophage reprogramming in therapeutic interventions.

Therapeutic approaches aimed at re-educating tumor-associated macrophages into immunostimulatory cells ^58^ may involve targeting the macrophage PI3Kγ intracellular signaling pathway ^59^, which inhibits NFκB activation through Akt and mTOR ^60^, thereby inducing an anti-inflammatory program that promotes immune suppression. Indeed, macrophage PI3Kγ inactivation stimulates and prolongs NFκB activation, promoting an immunostimulatory transcriptional program ^60^. The ability of PI3Kγ to toggle between immune stimulation and suppression aligns with our findings on GSK3 inhibition, as PI3Kγ inactivates GSK3 via phosphorylation of Ser^9^ (GSK3β) and Ser^21^ (GSK3α) ^30^. Threfore, GSK3 inhibition may contribute to the anti-inflammatory and immunosuppressive effects of PI3Kγ inactivation by enhancing MAFB expression. Moreover, MAFB might even participate in the macrophage reprogramming consequences of mTOR modulation ^61–63^ due to the known reciprocal interactions between GSK3 and mTOR ^64^. Hence, since the phosporylation of MAFB by GSK3 requires of a prior “priming” phosphorylation event ^30^, identification of the MAFB priming kinase(s) emerges as a potentially relevant effort in the search for additional targets for macrophage re-programming.

The significant impact of GSK3 inhibition by CHIR-99021 on the macrophage transcriptional profile mirrors the effects observed with other GSK3 inhibitors on the differentiation of monocyte-derived dendritic cells (MDDC) ^65^. Specifically, GSK3 inhibitors like LiCl, SB415286, or SB216763 redirect the GM-CSF+IL-4-dependent development of MDDC towards the generation of macrophage-like cells ^65^. Considering that MAFB expression is higher in monocyte-derived macrophages than in dendritic cells ^66^, it is reasonable to infer that GSK3 inhibitors also boost MAFB expression in cells differentiated in the presence of GM-CSF+IL-4, thereby supporting the modulation of GSK3 activity as a tool to alter lineage determination in myeloid cells.

The re-programming effect of GSK3 inhibition on *ex vivo* isolated alveolar macrophages bears significant therapeutic implications, particularly in severe COVID-19 cases where profibrotic monocyte-derived macrophages replace GM-CSF-dependent tissue-resident alveolar macrophages ^23,24,67^. Our observations reveal that exposure of *ex vivo* isolated alveolar macrophages to CHIR-99021 leads to the loss of the alveolar macrophage GM-CSF-dependent gene profile and the acquisition of a MAFB-dependent pro-fibrotic transcriptional landscape resembling that of M-CSF-dependent monocyte-derived macrophages. Given that MAFB serves as a negative regulator of GM-CSF signaling ^68^, downregulation of MAFB expression through modulation of GSK3 activity emerges as a rationale strategy to re-direct pathogenic monocyte-derived macrophages towards the acquisition of a lung-resident alveolar macrophage-like profile during severe COVID-19. This approach aligns with the proposed therapeutic use of GM-CSF in COVID-19 ^69,70^, as GM-CSF would impede or limit the MAFB-dependent upregulation of profibrotic and neutrophil-attracting factors observed in severe COVID-19 ^52^.

## ACKNOWLEDGEMENTS

This work was supported by grant PID2020-114323RB-I00 from Ministerio de Ciencia e Innovación to ALC, Dirección General de Innovación e Investigación Tecnológica de la Comunidad de Madrid (RETARACOVID, P2022/BMD-7274) to ALC, APK, PS-M and RD, Instituto de Salud Carlos III (Grant PI23/00224 to APK, Grant PI21/00989 to RD), Red de Enfermedades Inflamatorias (RICORS RD21/0002/0034) from Instituto de Salud Carlos III and cofinanced by the European Regional Development Fund “A way to achieve Europe” (ERDF) and PRTR to APK, and PID2021-123167OB-I00 from Ministerio de Ciencia, Innovación y Universidades and CSIC Talent Attraction program (20222AT010) to EO. This research work was also funded by the European Commission – NextGenerationEU (Regulation EU 2020/2094), through CSIC’s Global Health Platform (PTI Salud Global). AN-V was funded by YEI program contract from Comunidad de Madrid (PEJ-2021-AI/BMD-23327). IR was funded by a Formación de Personal Investigador predoctoral fellowship from Ministerio de Ciencia e Innovación (Grant PRE2021-097080). Figures were created with <u>Biorender.com</u>.

## AUTHORSHIP CONTRIBUTIONS

IR, CH, MT-T, BL-N, MT-S, FD-C, AN-V, RS and YS-P performed research and analyzed data; EO, PS-M, RD, AP-K and ALC designed research and analyzed data; FR-S and RG-L identified suitable patients for broncho-alveolar lavage; IR, APK and ALC wrote the paper.

## DISCLOSURE OF CONFLICTS OF INTEREST

The authors declare no competing financial interests.

## AVAILABILITY OF DATA AND MATERIALS

The dataset supporting the conclusions of this article is available in the Gene Expression Omnibus repository (http://www.ncbi.nlm.nih.gov/geo/) under accession GSE256538 (monocytes exposed to CHIR99021 or DMSO), GSE256208 (GM-MØ exposed to CHIR99021 or DMSO), GSE262463 (alveolar macrophages exposed to CHIR99021 or DMSO) and GSE266236 (GM-MØ after GSK3α/β knockdown).

## SUPPLEMENTARY FIGURE LEGENDS

**Figure S1.**
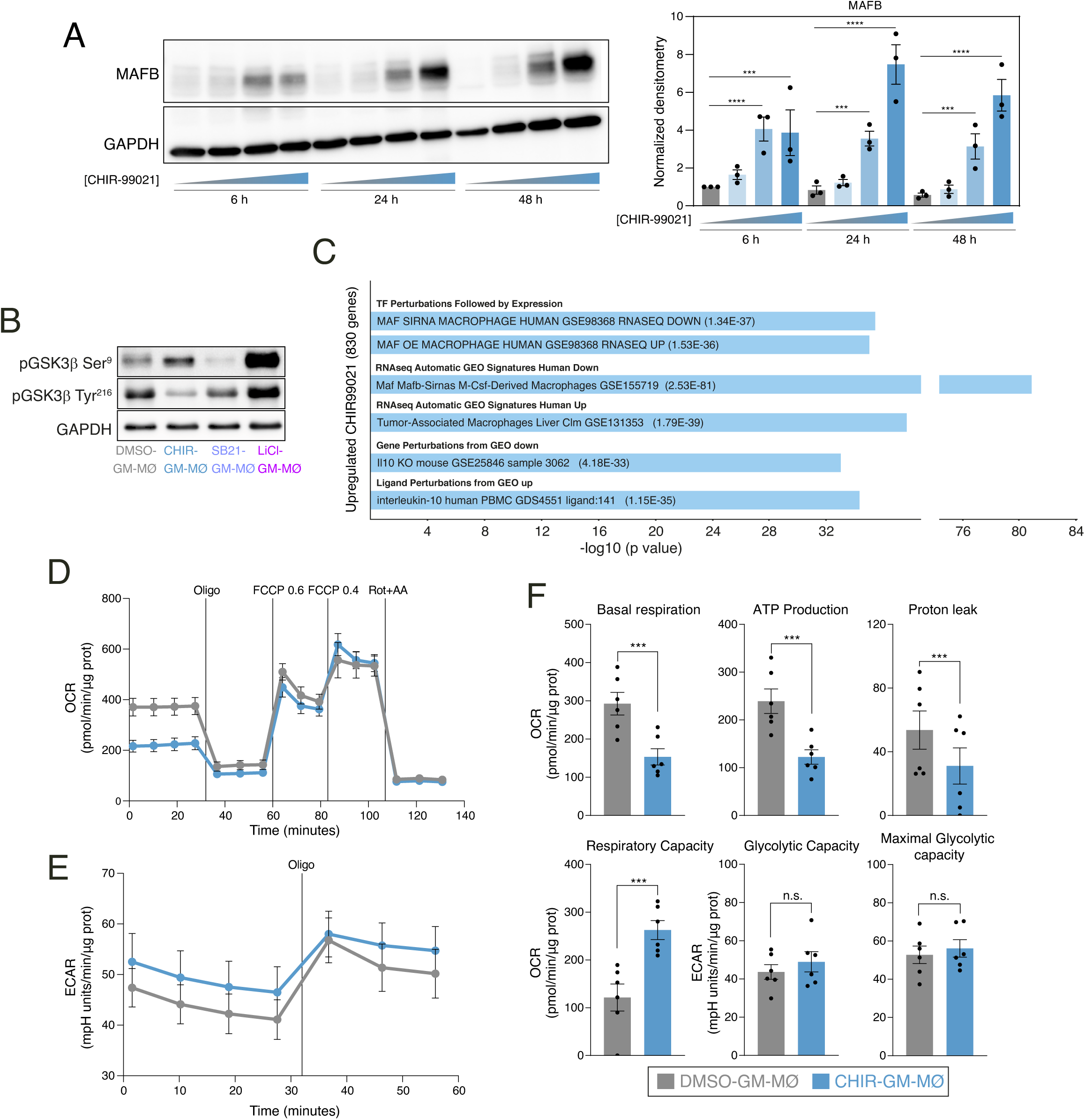
**A.** MAFB protein levels in GM-MØ exposed to DMSO or distinct CHIR-99021 concentrations (0.1μM, 1μM, 10μM) for 6-48 hours, as determined by Western blot (left panel). GAPDH protein levels were determined as protein loading control. Mean ± SEM of the MAFB/GAPDH protein ratio from three independent experiments are shown (right panel) (one-way ANOVA with Fisher LSD test: ***, p⍰<⍰0.005; ****, p<0.001). **B**. p-Ser^9^-GSK3β and p-Tyr^216^– GSK3β levels in GM-MØ treated with DMSO, CHIR-99021 (10 μM), SB-216763 (10 μM) or LiCl (10 mM) for 48 hours, as determined by Western blot. A representative experiment of three independent experiments is shown in the left panel. GAPDH protein levels were determined as protein loading control. **C**. Gene ontology analysis of the genes upregulated (|log2FC|>1; adjp<0.05) in CHIR-GM-MØ (relative to DMSO GM-MØ) using Enrichr and the indicated databases. **D**. Oxygen Consumption rate (OCR) profile of DMSO-GM-MØ and CHIR-GM-MØ monitored using the Seahorse Biosciences extracellular flux analyzer at the indicated time points. Cells were treated sequentially, as indicated, with 1 mM oligomycin (Oligo), 0.6 plus 0.4 mM FCCP, and 1 mM rotenone plus 1 mM antimycin A (Rot/AA). **E**. Extracellular acidification rate (ECAR), a proxy for the rate of lactate production, measured in DMSO-GM-MØ and CHIR-GM-MØ under basal conditions and after the stimulation with 1 mM oligomycin and at the indicated time points. **F**. Metabolic parameters obtained from the OCR and ECAR profiling after subtraction of the rotenone/antimycin-insensitive respiration. Basal OCR is the oxygen consumption rate in the absence of effectors, ATP turnover is considered as the oligomycin-sensitive respiration, maximal respiration is the OCR value in the presence of the uncoupler FCCP, glycolytic capacity is the ECAR value after the inhibition with oligomycin of the mitochondrial ATP synthesis (paired Student’s t test: ***, p<0.005; n.s., not significant). In **D-F**, results are normalized according to protein concentrations and presented as mean ± SEM of six independent samples.

**Figure S2.**
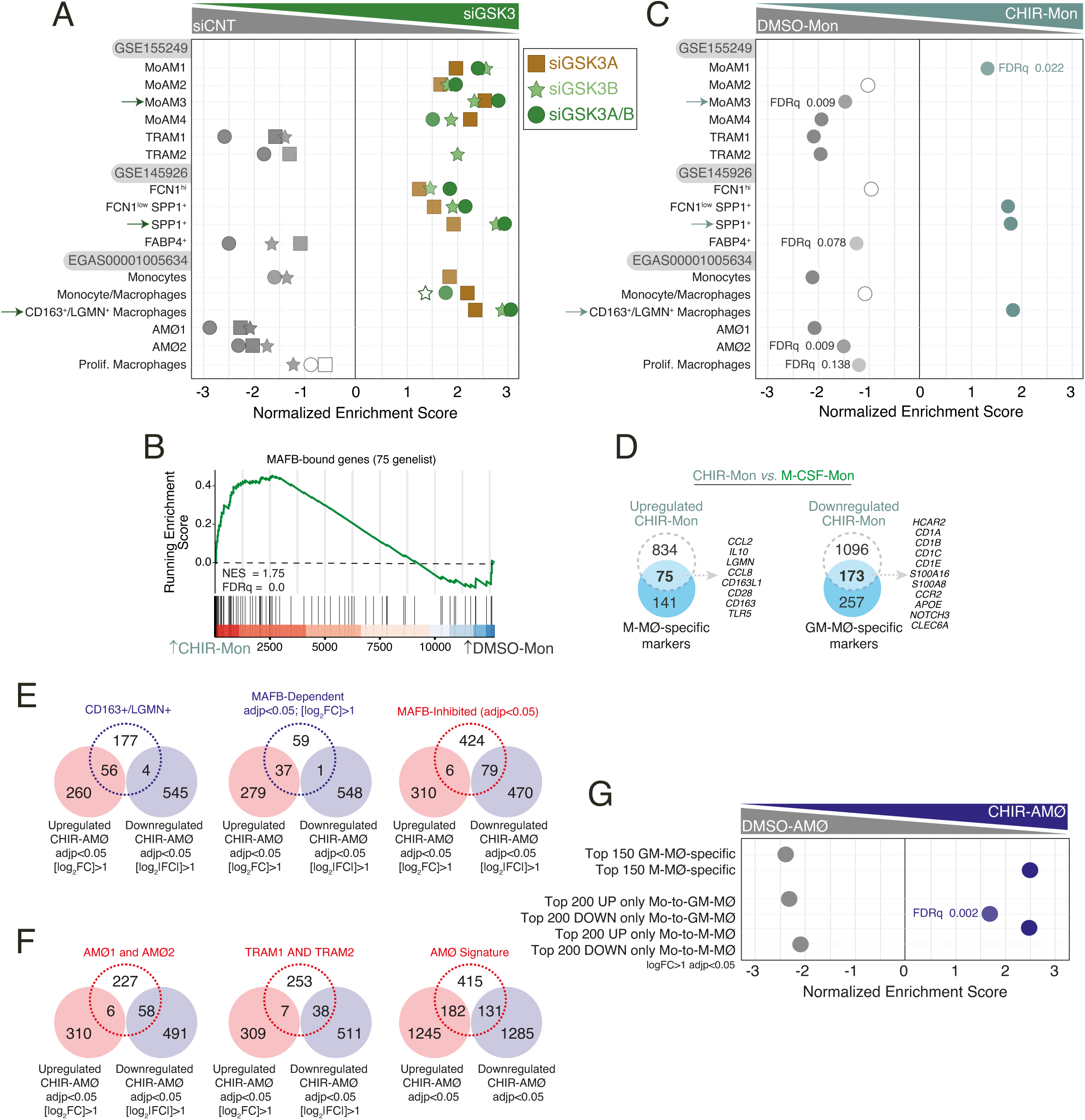
**A**. Summary of GSEA of the gene sets that characterize the macrophage subsets identified in severe COVID-19 ^23–25^ on the ranked comparison of siGSK3A-GM-MØ vs. siCNT-GM-MØ (square symbols), siGSK3B-GM-MØ vs. siCNT-GM-MØ (star symbols) and siGSK3A/B-GM-MØ vs. siCNT-GM-MØ. FDRq values are indicated only if FDRq>0.0; empty dots, Not Significant. **B.** GSEA of *bona fide* MAFB-bound genes (“75-genelist”) ^52^ on the ranked comparison of the transcriptomes of CHIR-Mon vs DMSO-Mon. NES and FDRq value is indicated. **C**. Summary of GSEA of the gene sets that characterize the macrophage subsets identified in severe COVID-19 ^23–25^ on the ranked comparison of the transcriptomes of CHIR-Mon vs. DMSO-Mon. FDRq values are indicated only if FDRq>0.0; empty dots, Not Significant. **D**. Overlap between the genes upregulated (|log2FC|>1; adjp<0.05) in CHIR-Mon (relative to M-CSF-Mon) and M-MØ-specific marker genes (left panel) and genes downregulated (|log2FC|>1; adjp<0.05) in CHIR-Mon (relative to M-CSF-Mon) and GM-MØ-specific marker genes (right panel), with indication of some shared genes. **E.** Overlap between the genes upregulated or downregulated in CHIR-AMØ (|log2FC|>1 and adjp<0.05) and the CD163+/LGMN+ pro-fibrotic macrophage cluster defined in EGAS00001005634 ^23^ (left panel), or the gene sets of MAFB-dependent (middle panel) and MAFB-inhibited (right panel) genes defined in GSE155719 ^52^. **F**. Overlap between the genes upregulated or downregulated in CHIR-AMØ (|log2FC|>1 and adjp<0.05, or just adjp<0.05) and the Alveolar Macrophage signature as defined in EGAS00001005634 (AMØ1 and AMØ2 clusters) ^23^ (left panel), GSE155249 (TRAM1 and TRAM2 clusters) ^25^ (middle panel) or in ^54^ (right panel). **G.** Summary of GSEA of the indicated gene sets (from GSE68061 and GSE188278) on the ranked comparison of the DMSO-AMØ and CHIR-AMØ transcriptomes. FDRq values are indicated only if FDRq>0.0.

**Figure S3.**
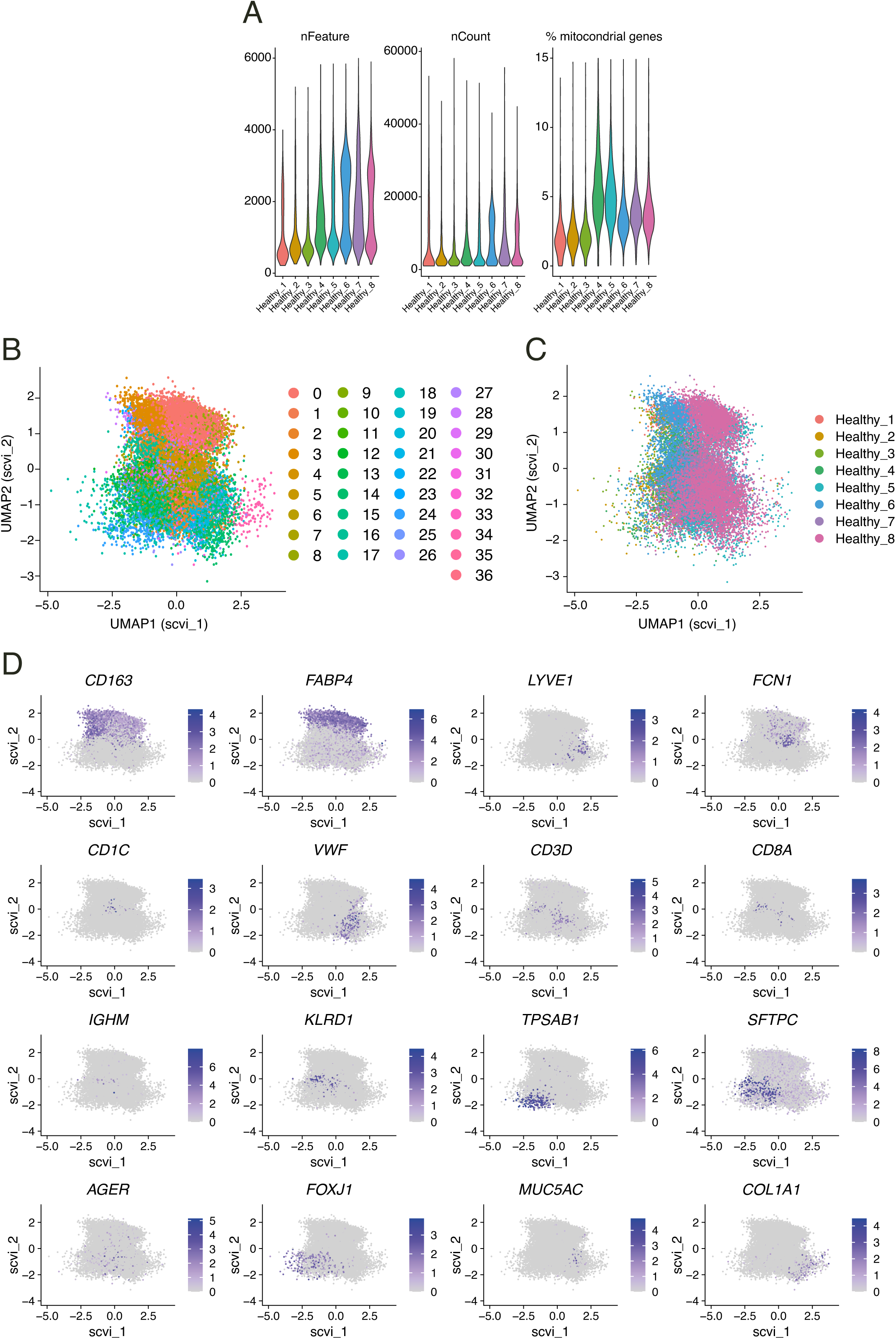
**A**. Number of cells from single cell-RNA sequencing on human lungs from eight independent donors (from GSE128033) ^47^, after filtering according to number of genes per cell (nFeature, > 200 and < 6000), Unique Molecular Identifiers (nCount, > 1000) and % of mitochondrial genes (< 15 %). **B**. UMAP (Uniform Manifold Approximation and Projection), with the scVI reduction, embedding of 21310 single-cell transcriptomes from human lungs as reported in GSE128033 ^47^, with color-coding indicating distinct cell clusters. **C**. UMAP (Uniform Manifold Approximation and Projection), with the scVI reduction, embedding of 21310 single-cell transcriptomes from human lungs as reported in GSE128033 ^47^, with color-coding indicating donor origin. **D**. Color-coded expression of the indicated genes after projection onto the UMAP embedding shown in **B-C**.

**Figure S4.**
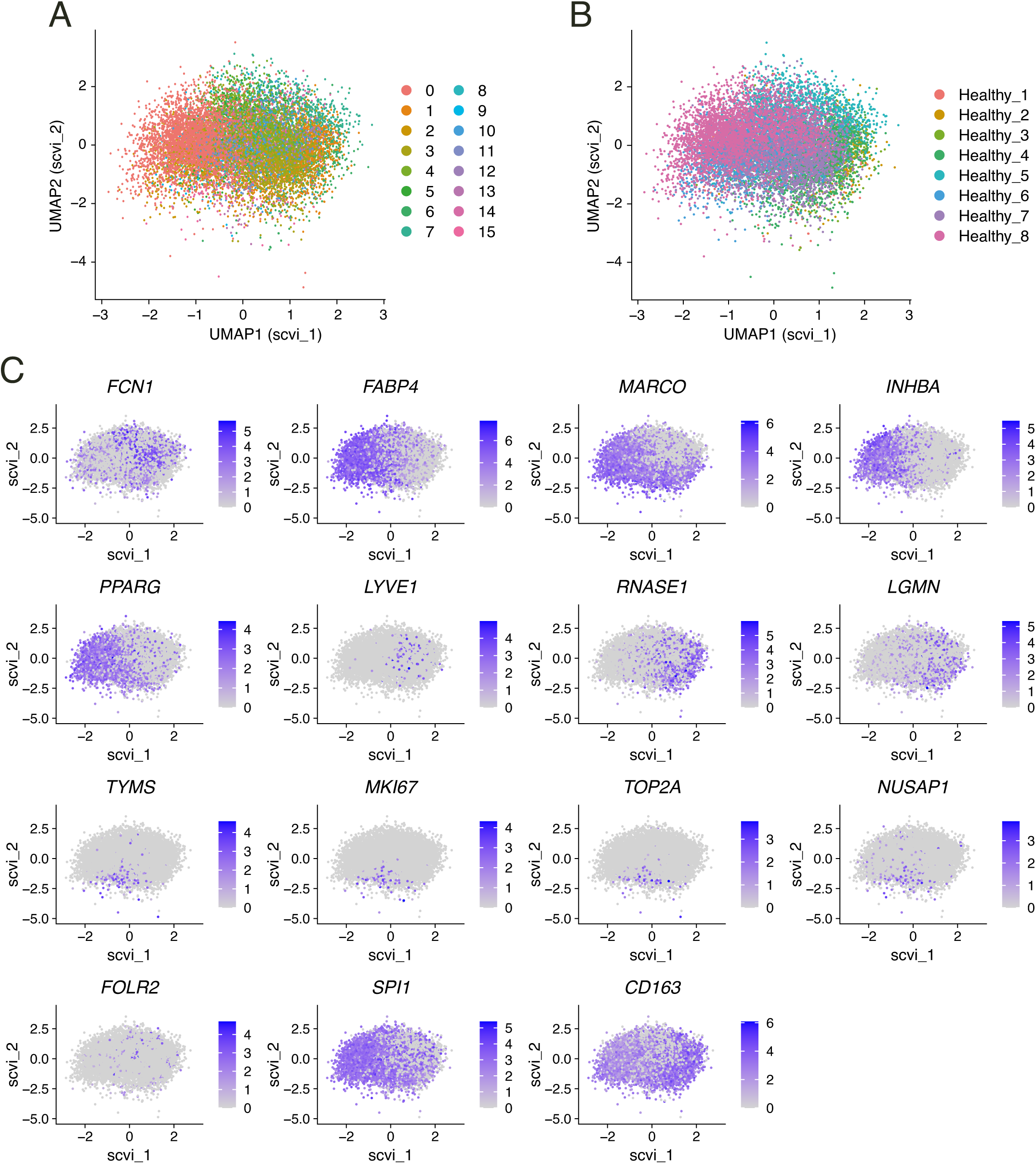
**A**. UMAP (Uniform Manifold Approximation and Projection), with the scVI reduction, embedding of single-cell transcriptomes from human lung macrophages as reported in GSE128033 ^47^, with color-coding indicating distinct cell clusters. **B**. UMAP (Uniform Manifold Approximation and Projection), with the scVI reduction, embedding of single-cell transcriptomes from human lungs as reported in GSE128033 ^47^, with color-coding indicating donor origin. **C**. Color-coded expression of the genes used for macrophage definition (*CD163*, *FABP4*, *LYVE1, FCN1*), as well as proliferation-associated genes (*TYMS*, *MKI67*, *TOP2A*, *NUSAP1*) and other *bona fide* macrophage marker genes (*SPI1*, *FOLR2*), after projection onto the UMAP embedding shown in **B-C**.

## SUPPLEMENTARY FIGURE LEGENDS

**Table S1.**
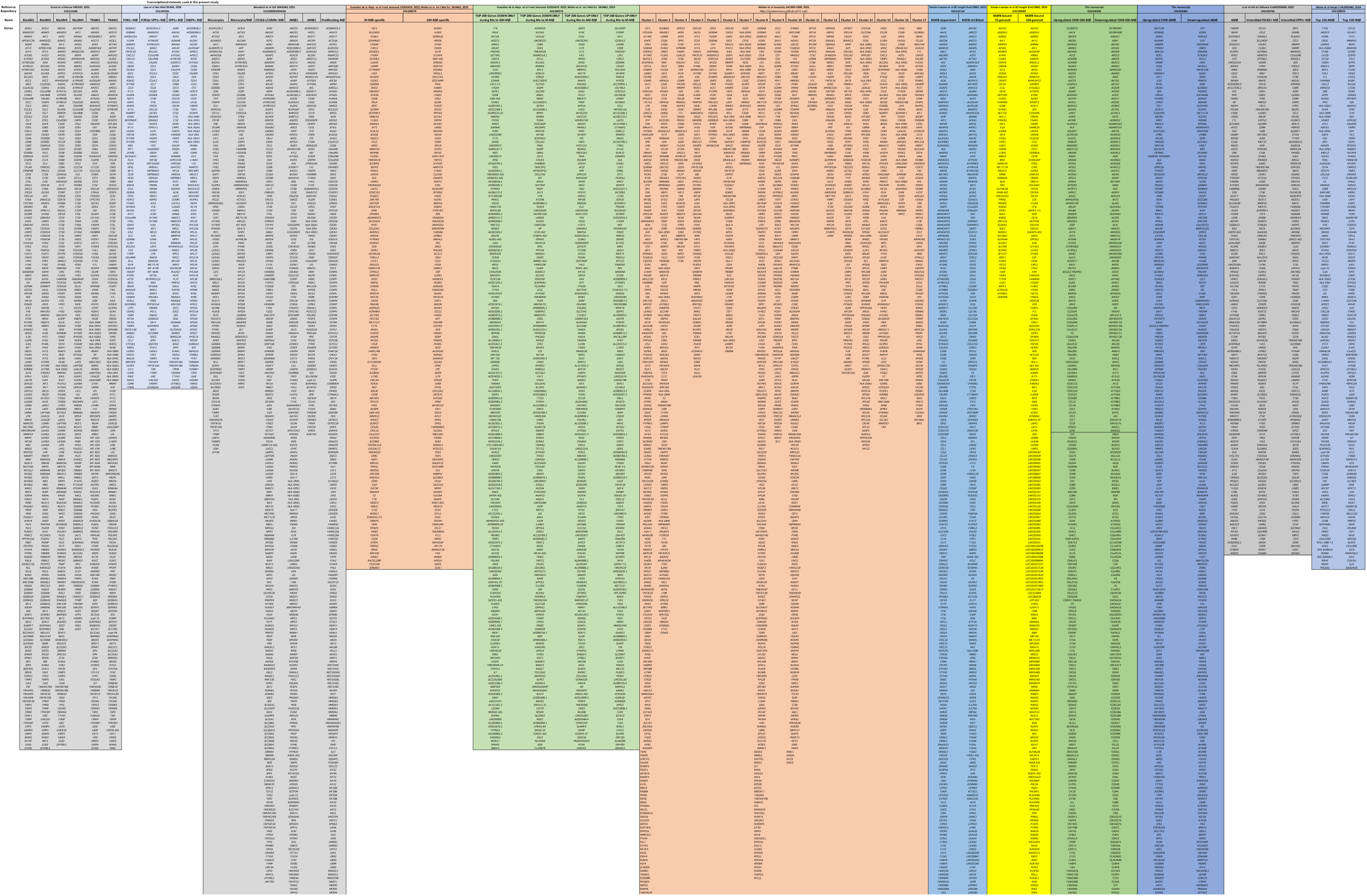
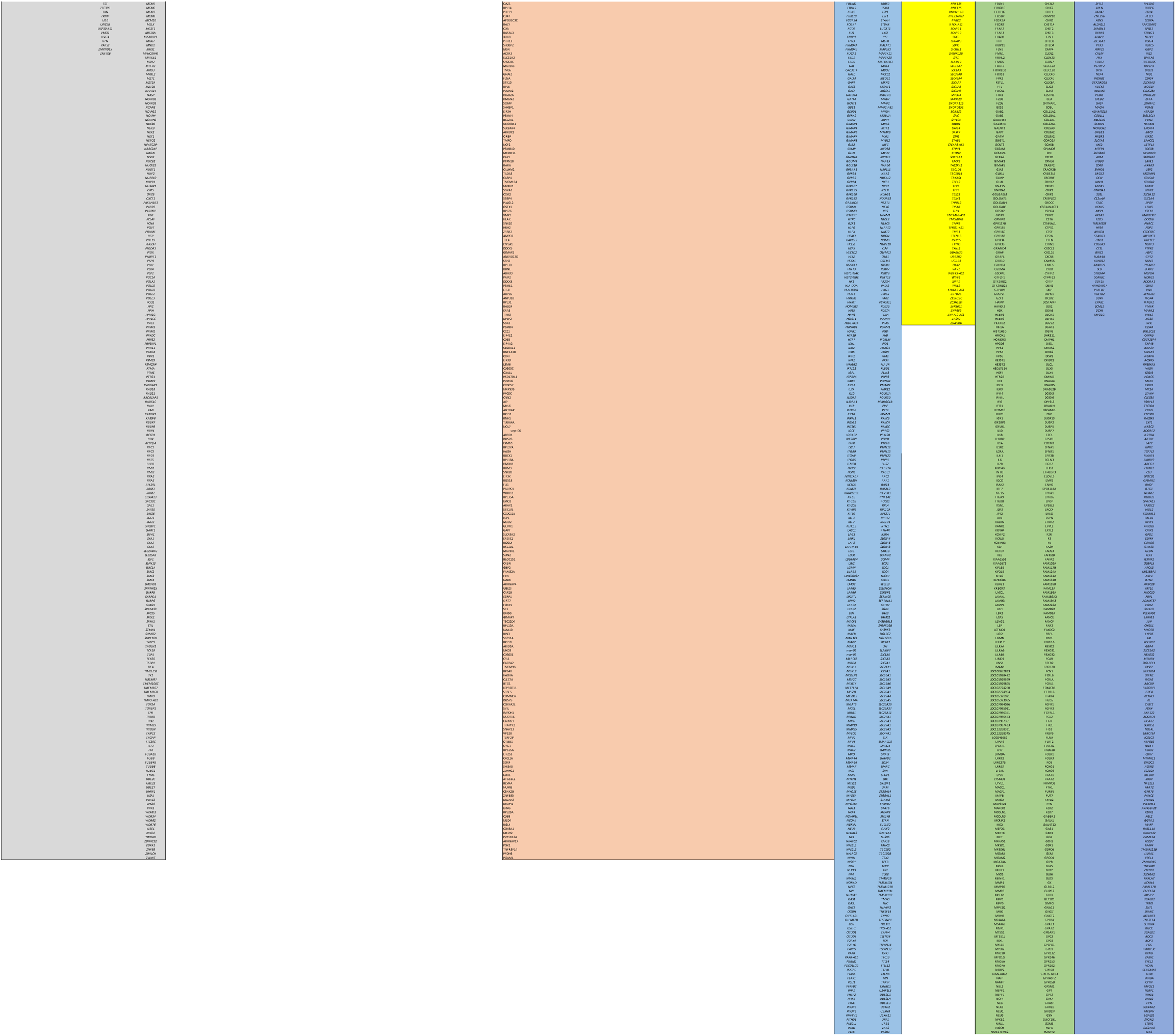
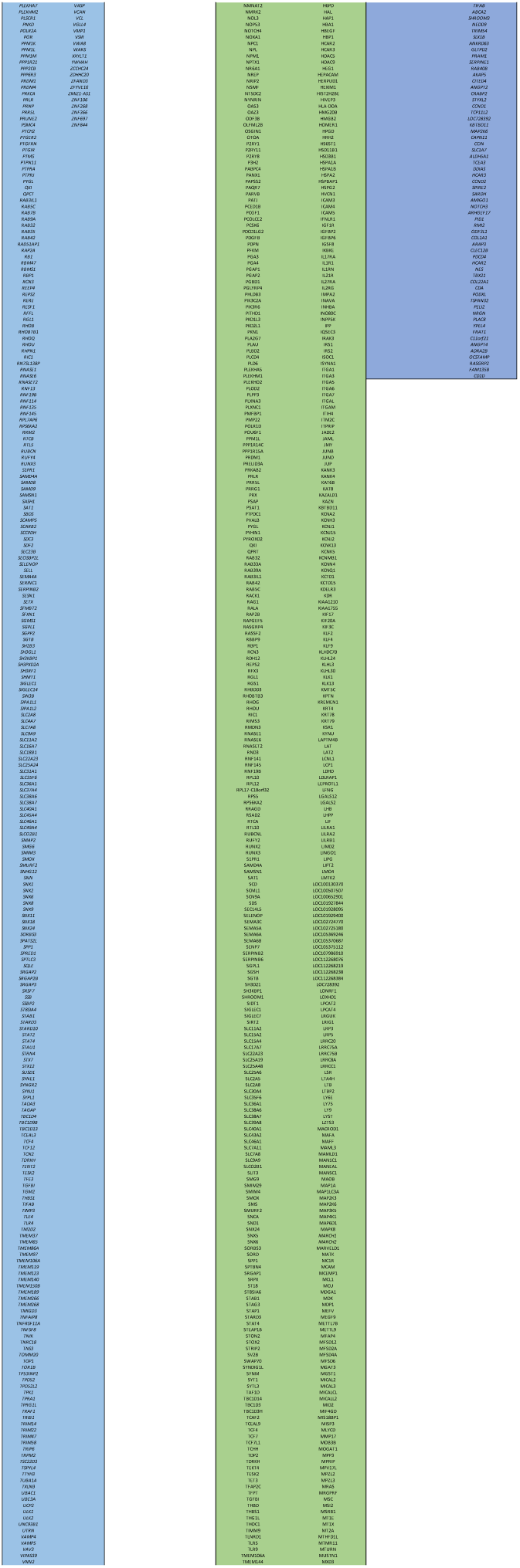
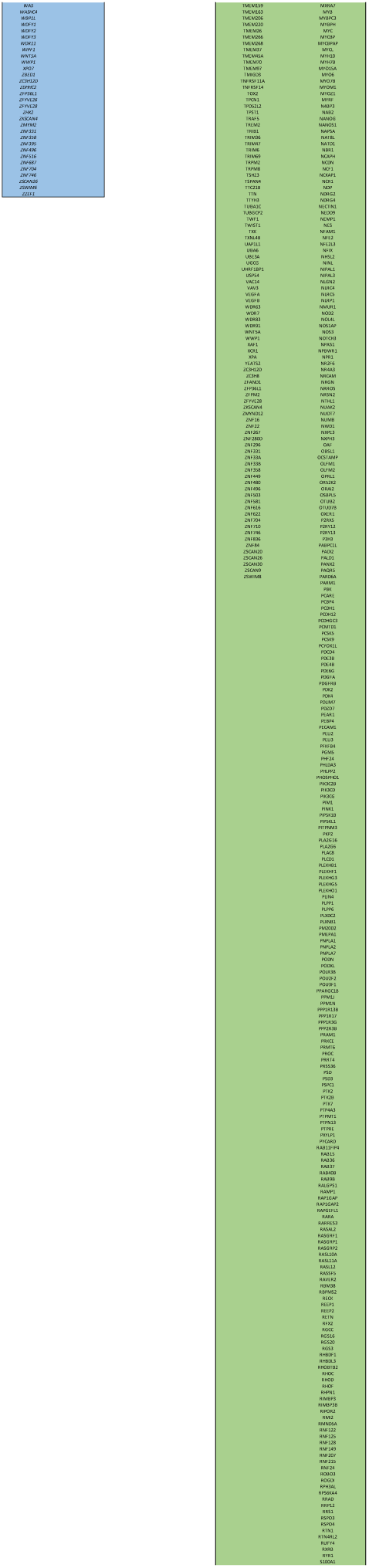
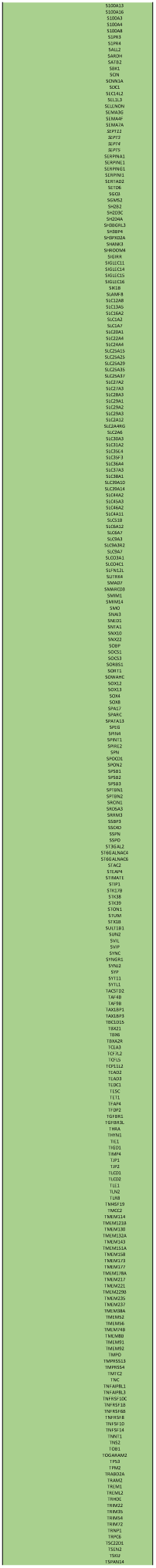
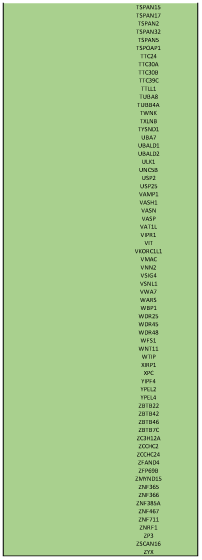
Transcriptional datasets used in the present study.

## Notes

### Competing Interest Statement

The authors have declared no competing interest.

### Summary of Updates

New Supplementary Figure 1A with kinetic and dose-response analysis; New Figure 7A-B, Supplementary Figure 3 and Supplementary Figure 4 to replaced all UMAP plots and to include Qc plots; wrongly formatted references throughout the manuscript have been corrected.

